# EPLINα controls integrin recycling from Rab21 endosomes to drive breast cancer cell migration

**DOI:** 10.1101/2024.06.27.600789

**Authors:** Niklas Z. Jäntti, Paulina Moreno-Layseca, Megan R. Chastney, Michal Dibus, James R. W. Conway, Veli-Matti Leppänen, Hellyeh Hamidi, Kathrin Eylmann, Leticia Oliveira-Ferrer, Stefan Veltel, Johanna Ivaska

**Affiliations:** Turku Bioscience Centre, University of Turku and Åbo Akademi University, FI-20520 Turku, Finland; University Medical Center Hamburg-Eppendorf (UKE), 20251 Hamburg, Germany; Hochschule Bremen, City University of Applied Sciences, 28199 Bremen, Germany; Department of Life Technologies, University of Turku, FI-20520 Turku, Finland; InFLAMES Research Flagship, University of Turku, 20014 Turku, Finland; Western Finnish Cancer Center (FICAN West), University of Turku, FI-20520 Turku, Finland; Foundation for the Finnish Cancer Institute, Tukholmankatu 8, FI-00014 Helsinki, Finland

**Keywords:** EPLINα, EPLIN isoforms, endosomal transport, integrin recycling, actin, Rab21, cell migration, breast cancer cells

## Abstract

EPLIN, an actin-binding protein, has been described as both a tumour promoter and tumour suppressor in different cancers. EPLIN isoform(α or β)-specific functions, which remain largely unknown, could explain these opposing roles. We observed distinct EPLIN-isoform localization; EPLINα is recruited to actin in plasma membrane ruffles and endosomes, while EPLINβ resides on actin stress fibers. We identified two EPLIN actin-binding regions and demonstrated EPLINα interaction with Rab21, an established regulator of β1-integrin endosomal traffic. EPLINα co-localizes with Rab21 and F-actin on recycling endosomes in an actin binding-dependent manner and supports β1-integrin recycling and cell migration. Using BioID, we identified coronin 1C as an EPLIN proximal protein, which localizes at Rab21-containing endosomes in an EPLINα-dependent manner. EPLINα expression was linked to increased breast cancer cell motility, and high EPLINα-to-EPLINβ ratio correlated with a mesenchymal phenotype in patient samples. Our work unveils unprecedented EPLIN isoform-specific functions relevant to breast cancer and beyond.

## Introduction

EPLIN (Epithelial Protein Lost in Neoplasm) is a LIM domain-containing, actin-binding protein. Since its discovery two decades ago, EPLIN has been linked to the regulation of cytoskeletal dynamics by bundling and crosslinking actin and implicated in cancer progression (1–3). EPLIN is encoded by the LIM domain and actin binding protein 1 (LIMA1) gene, but has two isoforms, α and β, generated by alternative promoters (4). Both isoforms are identical with the exception of an additional 160 amino acids at the N-terminus of EPLINβ. As its full protein name suggests, EPLIN has been described as a potential tumour suppressor due to its loss in epithelial tumours (1, 5). However, EPLIN has been shown to be upregulated in head and neck tumours, suggesting a pro-tumourigenic role (6, 7). EPLIN’s contradictory roles in cancer could be due to distinct or even opposing functions of its isoforms. However, this remains currently uninvestigated, highlighting the importance of studying the specific roles of each isoform. In addition to cytoskeletal functions, EPLIN is a structural component of adherens junctions, linking F-actin to the cadherin-catenin complex (8), participates in the accumulation of mitotic regulatory proteins at the cleavage furrow (9) and is regulated by kinases and phosphatases such as ERK (10) and CDC14A (11). EPLIN also binds to the endocytic molecule caveolin (12) and localizes at integrin adhesions in a PINCH-dependent manner in keratinocytes (13). PINCH is a part of the IPP complex (ILK-PINCH-parvin), which recruits molecules to integrin adhesion sites (14). Furthermore, EPLIN controls cholesterol absorption in the gut by aiding the recycling of NPC1L1, a transmembrane receptor expressed in the gastrointestinal tract (15). Despite these discoveries linking EPLIN to several important cell biological processes, the underlying molecular mechanisms and especially the specific roles of each EPLIN isoform remain largely unexplored. The endosomal transport of integrins, which allows internalization and delivery of receptors from and to the plasma membrane, is required for adequate integrin adhesion formation and disassembly during cell migration (16, 17). We have previously identified the actin-binding protein Swiprosin-1 (Swip1; EFHD2) as an interactor of the small GTPase Rab21 (18), and as an essential cargo-adaptor for endocytosis of active β1-integrins via the CLIC/GEEC pathway (19). Elevated integrin endocytosis by the Rab21-Swip1 axis in breast cancer cells increases cell migration, invasion and correlates with poor prognosis in triple-negative breast cancer (18). However, the mechanism regulating the recycling of the Swip1-Rab21 endocytosed integrins has not been investigated. Here, we demonstrate EPLINα as a direct interactor of Rab21 at recycling endosomes. We find that EPLINα, but not β, controls the recycling of integrins from these endosomes in an actin-dependent manner. EPLINα resides on Rab21-positive endosomes at localized puncta together with F-actin and coronin 1C, from where tubules are generated. Our study reveals a new function for EPLINα in integrin transport to drive migration of cancer cells, identifies isoform-specific proximity interactors and shows that the balance between EPLIN isoforms in cells correlates with their migratory potential.

## Results

### EPLINα interacts directly with Rab21 at endosomal compartments

Although our recent work has identified in detail the mechanism of Rab21-mediated recruitment and endocytosis of active integrins, there is a large gap in our understanding of key players that regulate the recycling of this cargo back to the plasma membrane. We therefore turned our attention to EPLIN, an actin-binding protein identified as a putative Rab21 interactor in our previous screen performed in MDA-MB-231 cells (18). We started by observing that MDA-MB-231 cells express both EPLIN isoforms, EPLINα and EPLINβ, with EPLINα being the predominant one (Fig. 1A-B). Immunofluorescence staining of EPLIN with an antibody recognizing both isoforms (referred to as EPLIN total from hereon) showed the endogenous protein localizing to actin-rich protrusions and stress fibers, but also on intracellular actin-positive endosomal-like structures in MDA-MB-231 cells (Fig. 1C). Co-expression of both EPLIN isoforms tagged to fluorescent proteins revealed shared and distinct subcellular localizations (Fig. 1D). Both isoforms localized to the actin cortex and lamellipodia, as previously reported (20, 21). However, EPLINβ was strongly enriched on actin stress fibers, while EPLINα was present in endosomal-looking structures. In order to investigate whether these structures corresponded to Rab21-containing endosomes, we performed super-resolution structured illumination microscopy (SIM) on cells transfected with mScarlet-EPLINα and Rab21-GFP and stained for F-actin. We observed that EPLINα localized to Rab21-positive endosomes in a punctate distribution around the endosome, which mirrored the localization of endosomal F-actin (Fig. 1E). To validate our observation, we performed Airyscan imaging to visualize endogenous EPLIN and Rab21. We found both proteins in endosomes containing active β1-integrin, which is a known cargo for Rab21-positive endosomes (18, 19, 22) (Fig. 1F). Line scan analysis of the intensity distribution of EPLIN, Rab21 and F-actin around the endosome showed that EPLIN and F-actin share the same punctate distribution around the endosome, while Rab21 is found in close vicinity to EPLIN, with partial overlap. Moreover, bimolecular fluorescence complementation (BiFC) experiments, where two proteins of interest are tagged with complementary halves of the Venus fluorescent protein (23), showed a positive interaction between the Rab21-EPLINα pair on endosomes decorated with F-actin (Fig. 1G). This interaction was absent in the mVenus control. In addition, microscale thermophoresis (MST) experiments demonstrated direct binding between GTP-analogue-loaded recombinant Rab21 and purified recombinant GST-EPLINα with sub-micromolar affinity (Fig. 1H). Together, these results indicate that EPLINα and Rab21 interact directly and localize at endosomes containing active β1-integrin, where EPLINα co-localizes with F-actin.

**Fig. 1.**
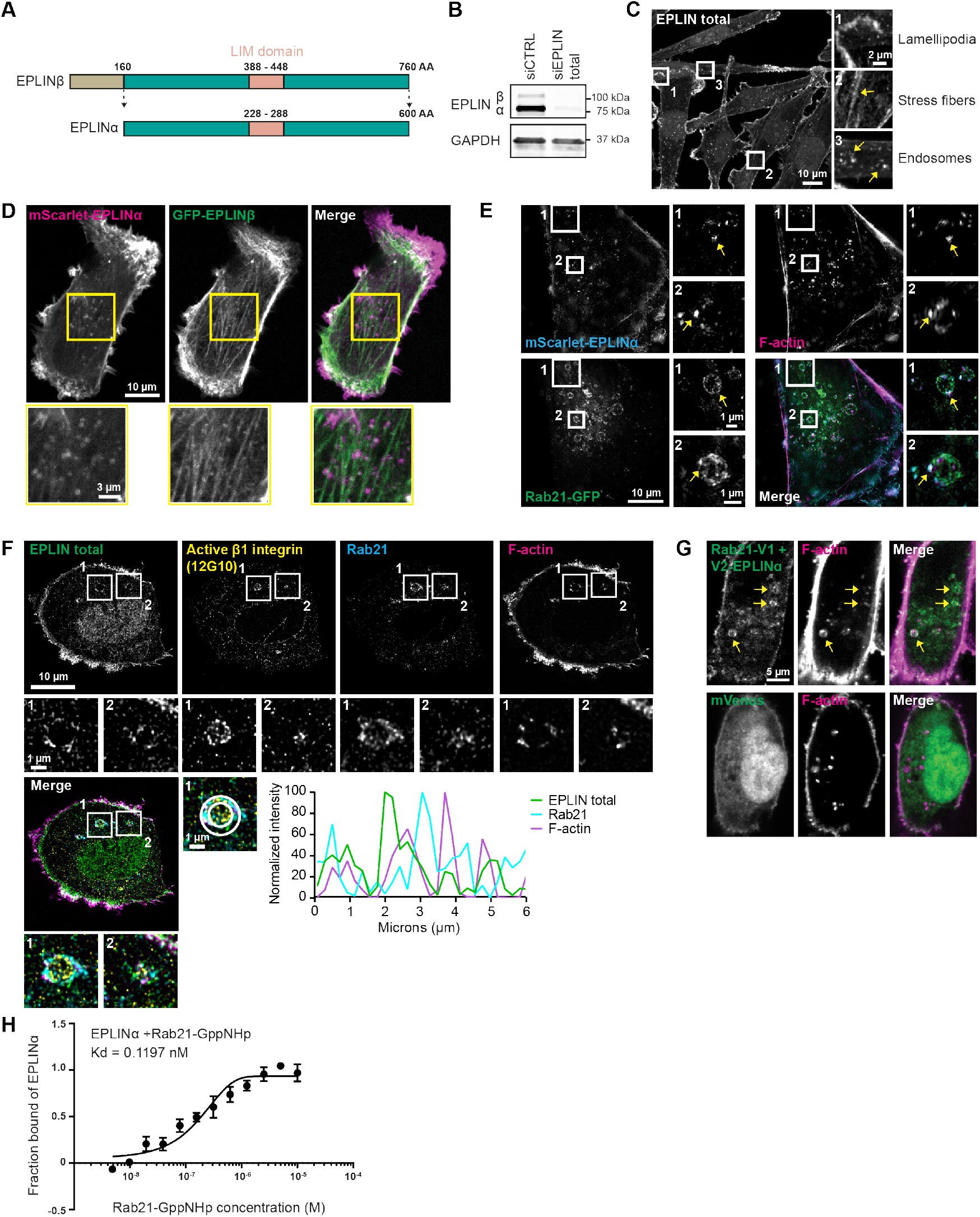
EPLINα interacts directly with Rab21 at endosomal compartments. A) Schematic representation of EPLIN isoforms. B) Representative immunoblot showing expression of EPLIN isoforms in MDA-MB-231 cells. Immunoblot is representative from 3 independent experiments. C) Representative MDA-MB-231 cells immunostained with antibodies that detect both EPLIN isoforms (total EPLIN). Insets highlight the different localizations of EPLIN: actin-rich protrusions (inset 1), stress fibers (inset 2) and endosomes (inset 3). Arrows point to structures of interest. D) MDA-MB-231 live cells overexpressing mScarlet-EPLINα and GFP-EPLINβ. The yellow square highlights a region of interest (ROI) showing localization of mScarlet-EPLINα primarily on endosomes and GFP-EPLINβ primarily on stress fibers. E) Representative SIM image of MDA-MB-231 cell expressing mScarlet-EPLINα and Rab21-GFP and labelled with Phalloidin-Atto 647N. Arrows point to EPLINα overlap with F-actin in Rab21-containing endosomes. F) Representative Airyscan image of maximum intensity projection of two slices from the middle of an MDA-MB-231 cell immunostained for total EPLIN, Rab21, active β1-integrin (12G10) and labelled with Phalloidin-Atto 647N. Insets (1, 2) show Rab21-positive endosomes containing active β1-integrin, EPLIN and F-actin. Histogram shows fluorescence intensity at the periphery of the endosome (indicated with a circle) to show colocalization between Rab21, EPLIN and F-actin. The raw Airyscan data was deconvolved using Huygens (Scientific Volume Imaging). G) Representative confocal microscopy BiFC images of MDA-MB-231 cells expressing the BiFC constructs Rab21-V1 and EPLINα-V2 or Venus alone as a control in cells stained with Phalloidin-Atto 647N. Arrows indicate BiFC colocalizing with F-actin. H) Binding of EPLINα to fluorescently labelled non-hydrolysable GTP analogue (GppNHp)-loaded Rab21 in microscale thermophoresis. Plot shows mean fraction bound and standard error of the mean from 3 independent experiments.

### EPLINα controls the recycling of active integrins

Prompted by our observation that EPLINα localized to Rab21- and active β1-integrin-containing endosomes, we asked whether it could play a role in the regulation of integrin trafficking. To answer this question, we performed a fluorescence microscopy-based β1-integrin internalization assay and measured the intracellular fluorescence intensity after internalization of cell surface-labeled active β1-integrins. Since EPLINβ is also expressed in MDA-MB-231 cells, we tested its potential contribution to integrin traffic regulation as well. Silencing of both EPLIN isoforms (Total Eplin) resulted in an increase of intracellular cell surface-labelled active β1-integrin after 15 minutes of internalization. Conversely, silencing only EPLINβ had no significant effect, suggesting that the EPLINα isoform specifically regulates active β1-integrin trafficking (Fig. 2A). This internalization assay was performed in the presence of serum, where the constant endocytosis of integrins is balanced with receptor recycling back to the plasma membrane (24). Thus, we hypothesized that the accumulation of active β1-integrin inside the cell could be caused by a defect in recycling. To study this in more detail, we performed another set of trafficking experiments, this time focusing on the recycling step by utilizing a previously established fluorescence quenching-based method (24). In this case, the internalization step was performed in the absence of serum to prevent recycling. As a result, the levels of endocytosed integrin were equal in the control and total EPLIN-silenced conditions, indicating that EPLIN is not regulating integrin endocytosis. Recycling was triggered by the addition of serum in a subsequent step, and the remaining signal inside the cell was quantified. More integrin signal remained in the EPLIN-silenced cells after this step, indicating a defect in recycling (Fig. 2B). Since our results showed a role for EPLINα in integrin recycling, we next aimed to investigate whether EPLINα localized at recycling endosomes. Rab11 mediates recycling of integrin cargo from perinuclear endosomes, while Rab4 mediates fast-recycling from early endosomes (16). Immunostaining of Rab11 in cells expressing endogenously (CRISPR-CAS9) tagged Rab21-mScarlet showed overlap of the two proteins on perinuclear endosomes, but not on smaller endosomes close to the plasma membrane (Fig. 2C). Moreover, endogenous EPLIN localized to endosomes positive for both Rab21-GFP and Rab11-mCherry (Suppl. Fig. 1B). We then used BiFC to visualize the Eplinα-Rab21 complexes in the cells and investigate their overlap with recycling markers Rab4, Rab11a and the Rab11-effector Rab11FIP5, which mediates the movement of cargo from early endosomes to Rab11-positive recycling compartments (25). EPLINα-Rab21 BiFC signal co-localized strongly with Rab11a and Rab11FIP5 and, to a lesser degree, with Rab4 (Fig. 2D, Suppl. Fig. 1C), indicating that EPLINα is mainly located at Rab11-positive recycling compartments. Together, these results support a possible role for EPLIN in the regulation of Rab21-bound integrin recycling via the Rab11 recycling pathway. Cargo recycling from endosomes requires the formation of tubules destined for the plasma membrane (26). Deconvolved Airyscan images of mScarlet-EPLINα and Rab21-GFP revealed Rab21- and EPLINα-positive tubules emanating from the endosomes (Fig. 2E). These data were further validated with live imaging of cells showing the generation of tubules from Rab21-containing endosomes arising from EPLINα puncta (Fig. 2F, Suppl. video 1). Together, these results indicate that EPLINα localizes to Rab21-positive recycling compartments, from where it mediates the recycling of active β1-integrin.

**Fig. 2.**
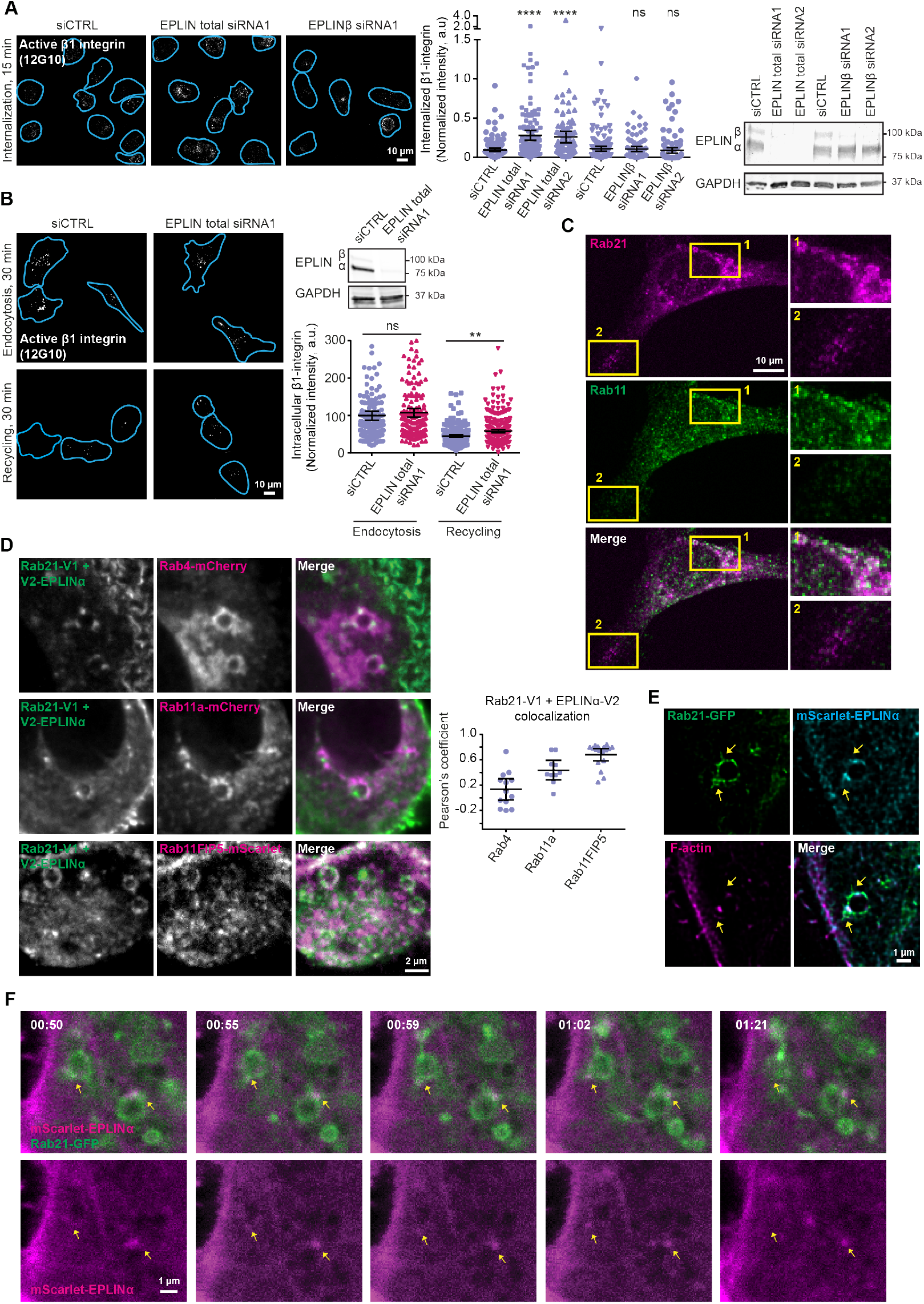
EPLINα controls the recycling of active integrins A) Schematic representation of EPLIN isoforms. A) Representative micrographs of active β1-integrin internalization at 15 minutes in control (siCTRL)- and EPLIN-silenced MDA-MB-231 cells (blue lines show the outlines of the cells defined by Phalloidin-Atto 647N labelling). Representative immunoblot to validate EPLIN silencing and quantification of the levels of internalized β1-integrin are shown. siCTRL, n = 142 cells; Total EPLIN siRNA1, n = 120 cells, Total EPLIN siRNA2, n= 102; siCTRL, n=177; EPLINβ siRNA1, n=95; EPLINβ siRNA2, n=107, from left to right. B) Representative micrographs of internalized active β1-integrin after 30 minutes of endocytosis and after 30 minutes of recycling. Representative immunoblot to validate EPLIN silencing and quantification of the levels of internalized β1-integrin are shown. For endocytosis, siCTRL, n =170 and EPLIN siRNA1, n=189 cells; for recycling, siCTRL, n = 198 and EPLIN siRNA1, n=246 cells. C) Representative images of MDA-MB-231 cells expressing endogenously CRISPR-CAS9 tagged Rab21-mScarlet immunostained for Rab11. Insets compare Rab11 signal on perinuclear and peripheral Rab21-positive endosomes. D) Representative BiFC images of MDA-MB-231 cells expressing the BiFC constructs Rab21-V1 and EPLINα-V2 and the indicated proteins fused to mcherry/Scarlet. Colocalization was quantified by measuring Pearson’s coefficients between BiFC and the indicated proteins at ROIs containing the endosomes in one cell. Rab4, n = 13; Rab11a, n=10 and Rab11FIP5, n=17 cells, from two independent experiments. E) Representative Airyscan images of MDA-MB-231 cells expressing Rab21-GFP, mScarlet-EPLINα and labelled with Phalloidin-Atto 647N. Arrows point to tubular structures emerging from the endosome. The raw Airyscan data was deconvolved using Huygens (Scientific Volume Imaging). F) Representative confocal images of live MDA-MB-231 cells expressing Rab21-GFP, mScarlet-EPLINα and labelled with SiR-actin. Arrows point to EPLIN puncta in Rab21-containing endosomes, from which tubules are generated. For A and B, scatter dot plots show data as the mean ± 95% Confidence Interval (CI). Statistical significance was assessed using Mann Whitney test for A and Kruskal–Wallis one-way analysis of variance (ANOVA) and Dunn’s post hoc test for B; n is the total number of cells pooled from three independent experiments. **P < 0.003; ****P < 0.0001; NS, not significant; a.u., arbitrary units.

### The actin binding function of EPLINα is required for endosomal localization and integrin recycling

EPLINα harbours at least two actin binding sites within the N-terminal and C-terminal domains flanking the centrally located LIM domain (20); however, their precise location remains unknown. Analysis of the primary sequence of EPLINα using ELM (Eukaryotic Linear Motif resource, http://elm.eu.org/) suggested a WH2 actin binding motif at the C-terminal part of the protein as one of the actin-binding sites. In addition, the structural prediction by AlphaFold3 (27) suggests that there is a short α-helix (107-121) in the N-terminal region that could contribute to actin binding (Fig. 3A). Therefore, we created three mutants of EPLINα lacking either the short N-terminal helix (ΔNHX, residues 107-121), the C-terminal WH2 motif (ΔWH2, residues 565-581) or both (ΔΔ) and used them to pulldown endogenous actin (Fig. 3B). We observed that, while each of the single mutants showed a partial decrease in actin binding, actin binding was almost completely abolished in the case of the double mutant (Fig. 3C-D). Moreover, the association between EPLINα WT and ARPC1B, a subunit of the ARP2/3 complex, suggests that both of the identified actin-binding sites are able to bind fibrillar actin (Fig. 3C, E). Live imaging of MDA-MB-231 cells co-transfected with mScarlet-EPLINα-WT and GFP-EPLINα-ΔΔ revealed that, while EPLINα-WT localized to endosomes, the endosomal localization of EPLINα-ΔΔ was decreased (Fig. 3F, Suppl. video 2). We then went on to test whether the loss of actin binding would affect the ability of EPLINα to promote recycling of active β1-integrin. Silencing of total EPLIN reduced the recycling of active β1-integrin, in line with Fig 2B. Re-expression of siRNA-resistant mScarlet-EPLINα-WT was sufficient to fully rescue the active β1-integrin recycling back to control levels, while expression of mScarlet-EPLINα-ΔΔ was unable to rescue the recycling defect (Fig. 3G). Thus, EPLINα requires its actin binding function to promote recycling of active β1-integrin.

**Fig. 3.**
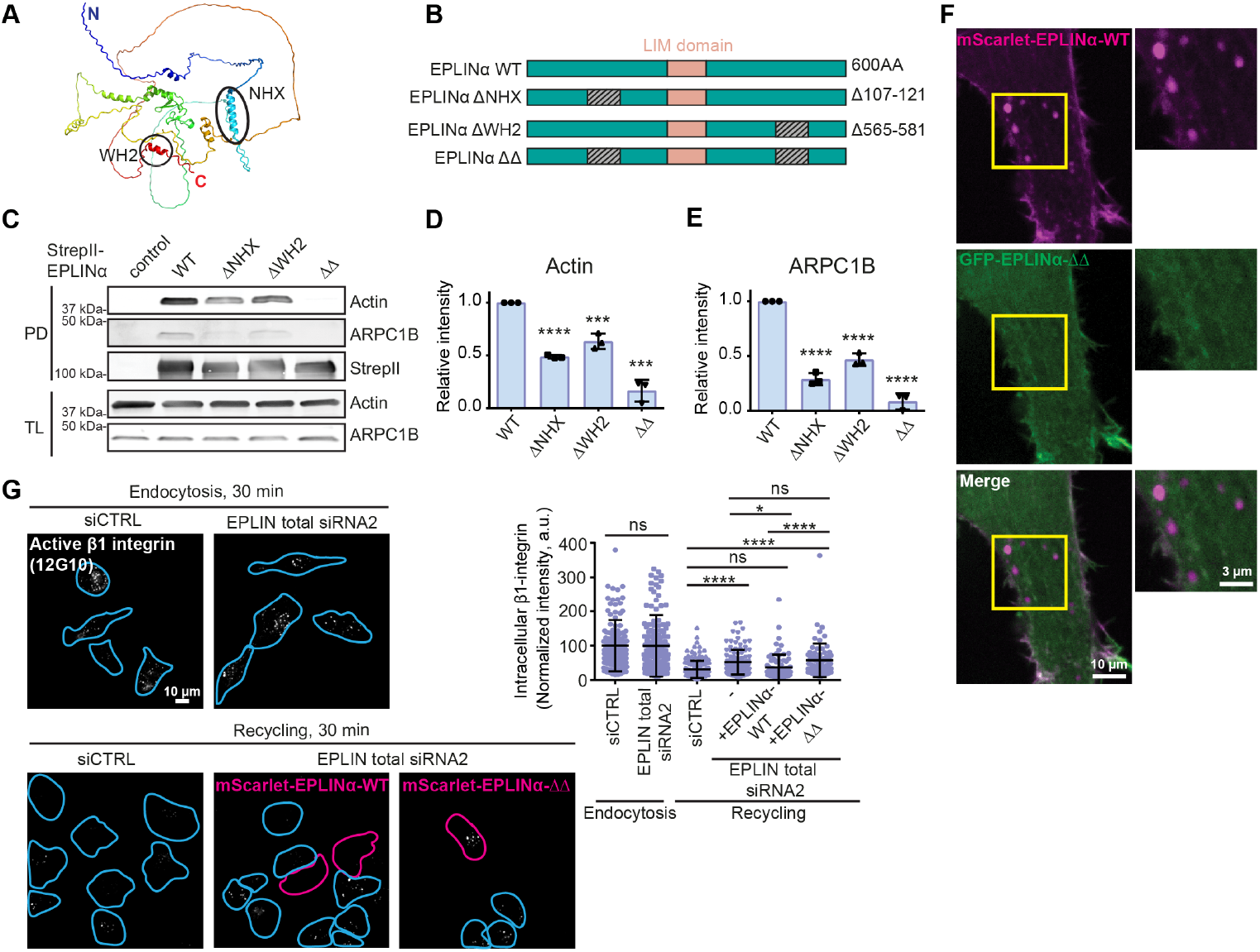
The actin binding function of EPLINα is required for endosomal localization and integrin recycling. A) AlphaFold prediction of EPLIN structure. The two actin-binding regions (N-terminal helix, NHX, and WH2 motif) are highlighted. B). Schematic representation of EPLINα-WT and the individual deletion mutants. C) Lysates of U2OS cells transfected with the individual StrepII-EPLINα variants were subjected to pulldown using Strep-Tactin and immunoblotted with indicated antibodies. D-E) Quantification of relative amounts of Actin and ARPC1B pulled down with individual StrepII-EPLINα variants in D. n=3 independent experiments. F) MDA-MB-231 live cells overexpressing mScarlet-EPLINα-WT and GFP-EPLINα-. ROI highlights endosomal localization of mScarlet-EPLINα-WT, but not of GFP-EPLINα-. G) Representative micrographs of internalized active β1-integrin after 30 minutes of endocytosis and after 30 minutes of recycling. Outlines of cells are shown in blue (siCTRL or Total EPLIN siRNA2) and magenta (cells expressing mScarlet-EPLINα-WT or -). Quantification of the levels of internalized β1-integrin are shown. For endocytosis, siCTRL, n=151 and EPLIN siRNA2, n=152; for recycling, siCTRL, n=148 and EPLIN siRNA2, n=105; EPLIN siRNA2 + EPLINα-WT, n=78 and EPLIN siRNA2 + EPLINα-, n=78 cells. Statistical significance for D and E was assessed using unpaired Student’s t test and for G using Kruskal–Wallis one-way analysis of variance (ANOVA) and Dunn’s post hoc test. For G, n is the total number of cells pooled from three independent experiments. *P<0.02; ***P<0.001; ****P<0.0001; NS, not significant; a.u., arbitrary units.

### BioID identifies proximity interactors for EPLINα and EPLINβ

Next we performed proximity biotinylation (BioID) coupled to mass spectrometry to identify proximity interactors for EPLINα and EPLINβ. BirA*-tagged EPLINα (BirA*-EPLINα), -EPLINβ (BirA*-EPLINβ), and BirA*-only control (BirA*) were stably expressed in two different breast cancer cell lines, MDA-MB-231 cells and HCC1937. In both cell types, the subcellular localization of BirA*-EPLINα and BirA*-EPLINβ was similar to endogenous/fluorescently tagged constructs, while BirA* showed no specific subcellular distribution and had a lower expression level (Fig. 4A, Suppl. Fig. 2A). To identify EPLIN proximity interactors, biotinylated proteins were isolated and analysed using mass spectrometry, and SAINTexpress (28) used to identify high-confidence bait-prey interactions, using BirA* as a negative control. A total of 91 proteins (between 54 and 57 per bait) were identified as proximity interactors (using BFDR of 0.05) of BirA*-EPLINα and BirA*-EPLINβ across both cell lines (Suppl. Table 1). Gene ontology (GO) analysis of the identified proteins revealed multiple actin and cell adhesion related terms, supporting the role of EPLIN in the regulation of actin and cell adhesion (29) (Fig. 4B and Suppl. Fig. 2B-C). Some proximity interactors (prey) were unique to EPLIN isoforms in specific cell lines and may represent isoform- and cell-specific interactors (Fig. 4C and Suppl. Fig. 2D). Many of these have established roles in actin dynamics and cell migration, such as LASP1 and lamellipodin (RAPH1) (30, 31) (group 5 in Fig. 4C), identified as EPLINα-specific interactors; and calponins CCN2 and CCN3 (32), identified as EPLINβ-specific interactors. Comparison of the enrichment of proximity interactors of EPLINα and EPLINβ in MDA-MB-231 cells revealed that even shared proximity interactors are differentially enriched by the two isoforms (Fig 4D). This could suggest that while EPLINα and EPLINβ share proximal proteins, there may be isoform-specific differences in their interactions (e.g. direct vs indirect, duration of interaction, distance between proteins, etc.). This could also mean that, given the different localizations of the isoforms in cells, each isoform could recruit the shared interactor to different cellular compartments. Notable identified proximity interactors with a potential role in the regulation of endosomal traffic include supervilin 1 (SVIL), cortactin (CTTN) and coronin 1C (CORO1C). Supervillin has been shown to bind EPLIN directly, and both proteins localize to the cleavage furrow in HeLa cells (9, 33). It has also been reported to localize to endosomes and to promote the recycling of β1- and β3-integrins (34). Cortactin is an activator of the Arp2/3 complex and has been implicated in the regulation of endosomal actin dynamics during cargo recycling (35). Coronin 1C (CORO1C) is an actin-binding protein that localises to both lamellipodia and the actin cortex, as well as to actin-regulated cargo domains on ER-related endosome buds, where it enables fission of sorting endosomes (35–38). Based on the literature and our findings, coronin 1C and EPLIN share a similar localization pattern in cells: coronin 1C localizes to the actin cortex and lamellipodia, where both EPLIN isoforms localize, and it also localizes to endosomes, where EPLINα is the predominant isoform, prompting us to explore coronin 1C in more detail.

**Fig. 4.**
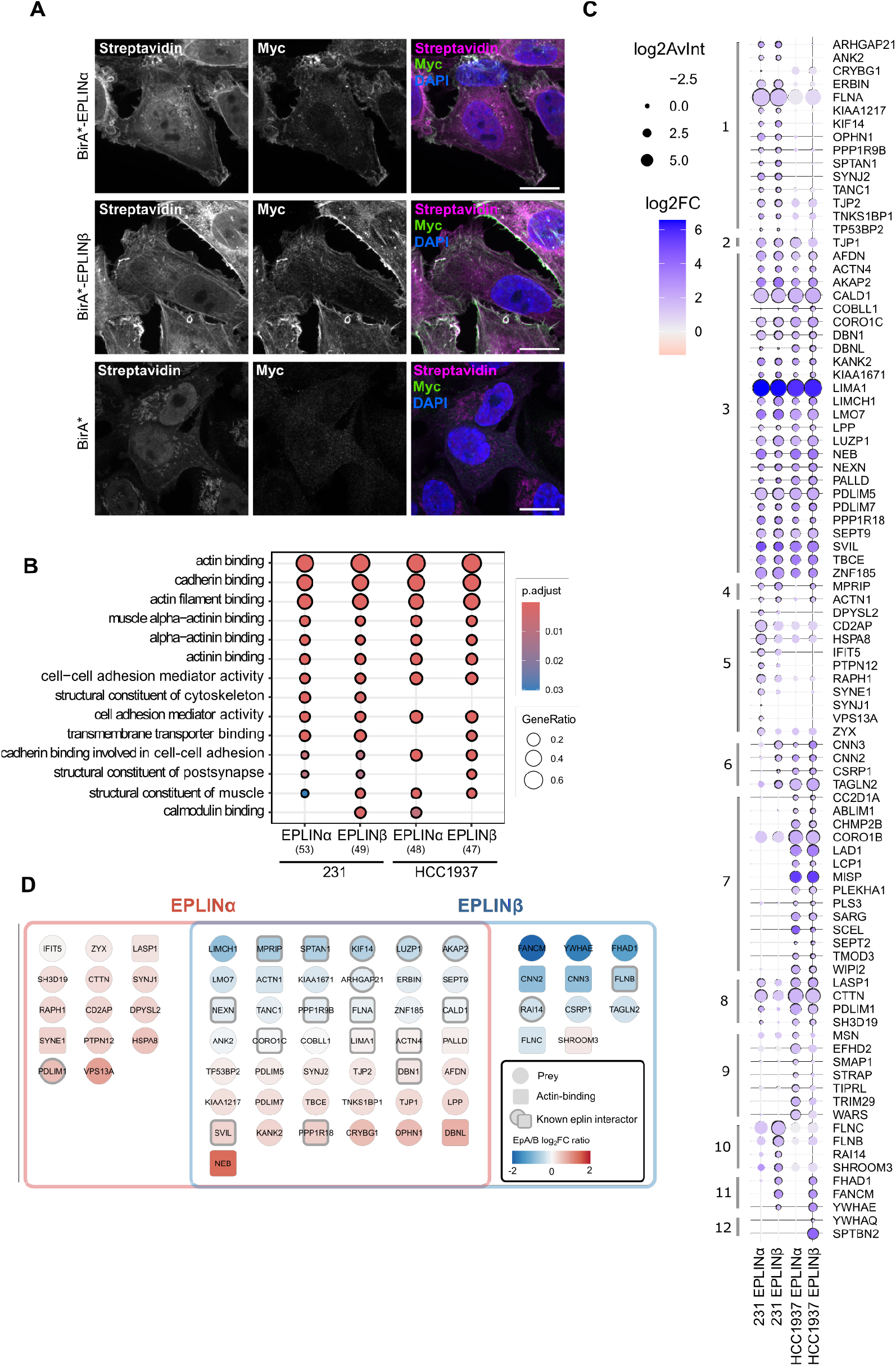
Identification of the EPLINα and EPLINβ proximity interactome by proximity biotinylation (BioID). A) Immunofluorescence images of MDA-MB-231 cells stably expressing BirA*-tagged EPLINα, EPLINβ or BirA* control. Biotin was added for 24 hours to induce labelling and biotinylated proteins detected by fluorescently-conjugated streptavidin. BirA*-tagged EPLIN was detected by anti-myc antibody and the nuclei were stained using DAPI. B) Gene ontology of prey identified by BioID shows multiple enriched terms relating to actin-binding/regulation (Molecular Function). Number of gene names used for analysis indicated in brackets. p.adjust; adjusted p value, GeneRatio; ratio of genes identified in terms. GO analysis performed using ClusterProfiler (39). C) Dotplot of high-confidence (BFDR 0.05) proximity interactors (prey) of BirA*-EPLINα and BirA*-EPLINβ in MDA-MB-231 and HCC1937 cells. Affinity-purified biotinylated proteins were analysed by mass spectrometry and high-confidence (BFDR 0.05) interactors identified using SAINTexpress (28). Dot size indicates the average intensity of three repeats (log2; log2AvInt), and colour indicates fold change over BirA* control (log2; log2FC). High-confidence (BFDR 0.05) interactions are indicated with a black outline. Prey are organised by hierarchical clustering. D) Unique and shared proximity interactors (BFDR 0.05) identified by MDA-MB-231 cells expressing BirA*-EPLINα and BirA*-EPLINβ. Colour indicates ratio between EPLINα and EPLINβ log2 fold change over BirA* only control (EPLINα log2 fold change/EPLINβ log2 fold change). Square nodes indicate actin-binding proteins (UniProt Annotated Keywords), and thick grey borders indicate known EPLIN interactors (BioGRID).

### Coronin 1C localizes at Rab21-containing endosomes in an EPLINα-dependent manner

The initiation of the early steps of endosome fission requires the WASH complex and activation of Arp2/3-mediated actin branching to drive membrane constriction (40). However, recent findings suggest that the halting of branching nucleation by Arp2/3 inhibition plays an important role in the later steps of fission, allowing access for other players in the fission machinery (41). One such Arp2/3 inhibitor that has been implicated in this process is coronin 1C. As EPLIN has also been shown to act as an inhibitor of Arp2/3 (20), we hypothesized that EPLINα and coronin 1C could work in conjunction to facilitate the inhibition of actin branching that is needed for fission of recycling endosomes to occur. Thus, we investigated whether coronin 1C could associate with EPLINα at Rab21-positive endosomes. SIM and Airyscan imaging of coronin 1C revealed that it localizes to Rab21-positive endosomes as distinct puncta overlapping with EPLINα and F-actin (Fig. 5A-B), concordant with our BioID results. Interestingly, coronin 1C cytoplasmic signal and localization on Rab21 endosomes was significantly reduced after EPLIN silencing (Fig. 5C), while the total protein levels of coronin 1C remained unchanged (Fig. 5D). This indicates that EPLINα is necessary for the recruitment of coronin 1C to endosomes.

**Fig. 5.**
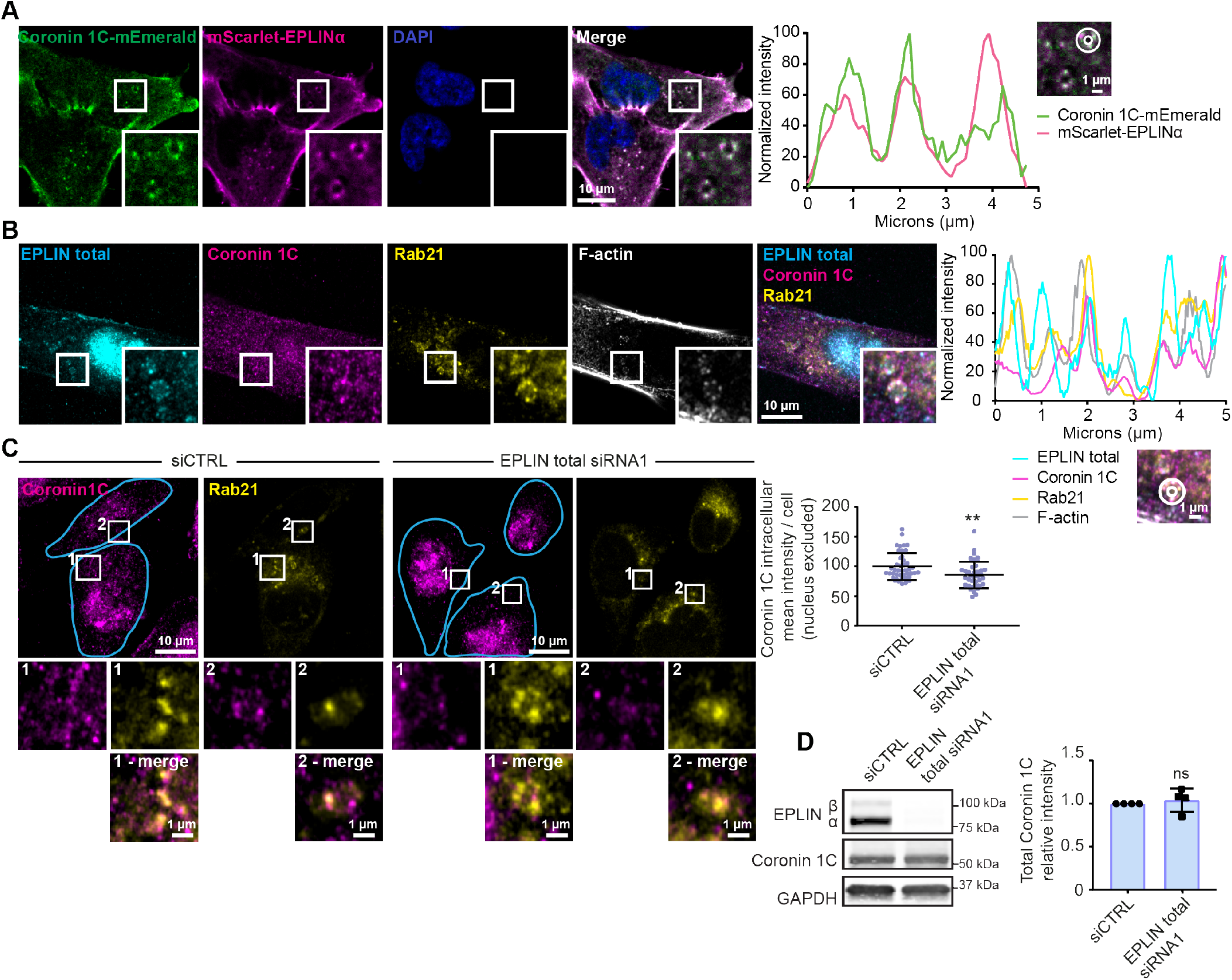
Coronin 1C localizes at Rab21-containing endosomes in an EPLIN-dependent manner. A) Representative confocal micrograph of a MDA-MB-231 cell expressing mScarlet-EPLINα and coronin 1C-mEmerald. Histogram shows fluorescence intensity at the periphery of the endosome (indicated with a circle) to show colocalization between EPLINα and coronin 1C. Cell nuclei were stained with DAPI. B) Representative Airyscan image of a MDA-MB-231 cell immunostained for total EPLIN, coronin 1C and Rab21, and labelled with Phalloidin-Atto 647N. Histogram shows fluorescence intensity at the periphery of the endosome (indicated with a circle) to show colocalization between Rab21, EPLIN total and coronin 1C. C) Representative images of control (siCTRL)- and total EPLIN-silenced MDA-MB-231 cells immunostained for coronin 1C and Rab21. Quantification of intracellular coronin 1C intensity, excluding nucleus, is shown. siCTRL n=43 cells and EPLIN siRNA1 n=37 cells. Cell outlines are shown in blue. D) Immunoblot and quantification of coronin 1C expression in control (siCTRL) and EPLIN-silenced (EPLIN total siRNA1) cells. n=4 independent experiments. Statistical significance was assessed for C using using two-sided Mann–Whitney test and for D using unpaired Student’s t test. For C, n is the total number of cells pooled from three independent experiments. **P<0.01; ns, not significant.

### High expression of EPLINα correlates with triple negative molecular subtype in breast tumours

Molecular regulators of integrin intracellular transport, including Rab21 and Swip1, have been shown to be over-expressed in clinical breast cancer samples (18, 42). To assess whether this was the case for EPLINα, we analysed the expression of EPLIN in a cohort of 105 human breast cancer samples by immunoblotting of tumour lysates (Fig. 6A). We chose this approach as the identical amino acid sequence of the two isoforms (apart from the β-specific N-terminus) precludes specific detection of EPLINα with the currently available antibodies. We found that the two EPLIN isoforms are expressed in different ratios among the tumours. Therefore, in order to interrogate the contribution of each isoform to the clinical outcome, we categorized them into four different groups according to the expression ratio: EPLINα high (α>β), EPLINβ high (β>α), equal expression of both isoforms (α=β) or no EPLIN expression. Our cohort had a similar percentage of tumours in each group (Fig. 6B). Among the ER-negative tumours, a high number were EPLINα high; and also a high number of the EPLINα high tumours belonged to the triple negative breast cancer subtype (Fig. 6C-D). This is interesting as our earlier work has shown that Rab21-Swip1-mediated integrin endocytosis and cell motility are linked to poor clinical outcome specifically in triple-negative breast cancer (18). Moreover, microarray data corresponding to the same breast cancer tissue cohort revealed that vimentin mRNA levels were significantly higher in the EPLINα high group compared to the other categories, further implying a more mesenchymal phenotype for tumours with high EPLINα expression (Fig. 6E). Altogether, these results show that breast tumours where EPLINα is the predominant isoform are more mesenchymal and mainly represent the triple-negative molecular subtype. Our observations also suggest that the ratio of expression of EPLIN isoforms, rather than overall expression, correlate with the tumour phenotype and may be important for the clinical outcome in breast cancer.

**Fig. 6.**
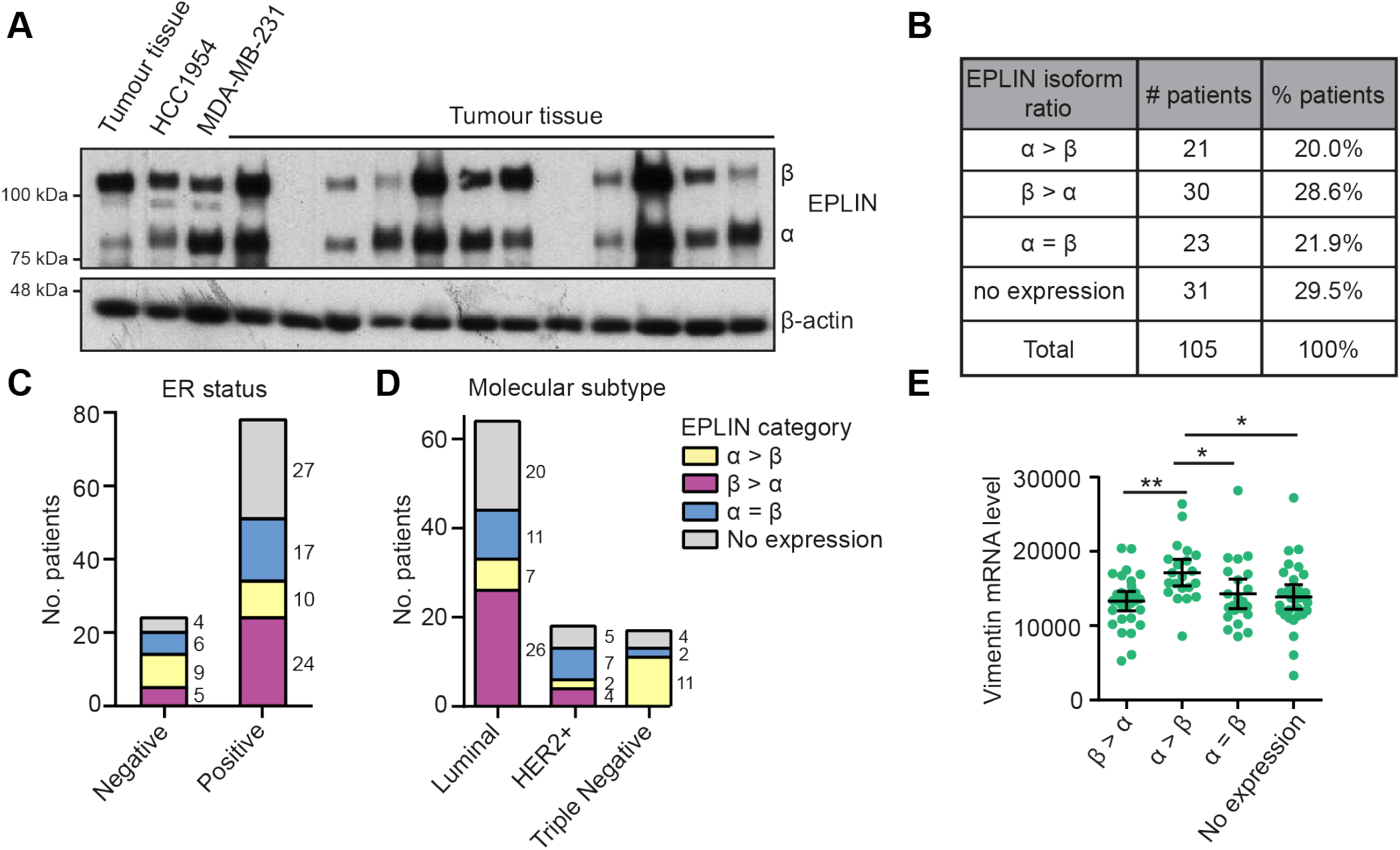
High expression of EPLINα correlates with triple negative molecular subtype in breast tumours. A) Representative immunoblot showing expression of EPLIN isoforms in 13 breast tumours from a cohort of 105 patients. Lysates of HCC1954 and MDA-MB-231 cells were used as controls. B) Tumours from 105 patients were categorized into four groups: EPLIN α>β, β>α, α=β or No expression according to the ratio of expression of EPLIN isoforms analysed by immunoblotting. Table shows the number and percentage of tumours in each category. C) ER status and D) molecular subtype of the tumours in each EPLIN category. E) Analysis of vimentin mRNA expression using qPCR in 105 breast tumours. Statistical significance was assessed using Kruskal–Wallis one-way analysis of variance (ANOVA) and Dunn’s post hoc test, *P< 0.04, **P= 0.0044.

### EPLINα mediates cell migration and correlates with a more mesenchymal phenotype in breast cancer cells

The heterogeneous expression of EPLIN isoforms in the clinical samples led us to hypothesize that cell lines where EPLINα is the predominantly expressed isoform would have higher motility and a more mesenchymal-like phenotype compared to cells where EPLINβ is predominant. To test this hypothesis, we used five different breast cancer cell lines: HCC1937, T47-D, MCF7, MDA-MB-231 and BT-549. Analysis of morphometric parameters gave high circularity and roundness scores for the cell lines HCC1937, T47-D and MCF7, whereas the cell lines MDA-MD-231 and BT-549 showed a significantly more elongated morphology (Fig. 7A-B), indicative of a less epithelial morphology. Immunoblotting of EPLIN showed that all cell lines with high circularity and roundness scores express predominantly EPLINβ. In contrast, the cell lines with a more elongated morphology express predominantly EPLINα (Fig. 7C). Moreover, the elongated, EPLINα-high cell lines are characterised by a high expression of vimentin and loss of E-cadherin (Fig. 7C), an expression pattern that is associated with a mesenchymal morphology and increased cell motility and invasiveness (43). This correlated with immunostaining of EPLINβ at cell-cell junctions in HCC1937 cells (Suppl. Fig. 3). Live cell microscopy and tracking of cell migration showed that the EPLINα-high MDA-MB-231 and BT-549 are markedly more motile compared to the cell lines predominantly expressing EPLINβ (Fig. 7D, Suppl. videos 3-7), in line with our hypothesis. To analyse EPLIN’s contribution to cell migration, we performed siRNA knockdown and rescue experiments in MDA-MB-231 cells. Total silencing of EPLIN significantly decreased migration speed, which was fully rescued back to control levels by expressing GFP-EPLINα-WT. Importantly, the actin binding-deficient mutant of EPLINα (GFP-EPLINα-ΔΔ) as well as GFP-EPLINβ-WT and GFP-EPLINβ-ΔΔ were all unable to rescue migration speed (Fig. 7E, Suppl. videos 8-13). Thus, EPLINα supports cell migration in MDA-MB-231 in an actin binding dependent manner, presumably through its ability to support integrin recycling.

**Fig. 7.**
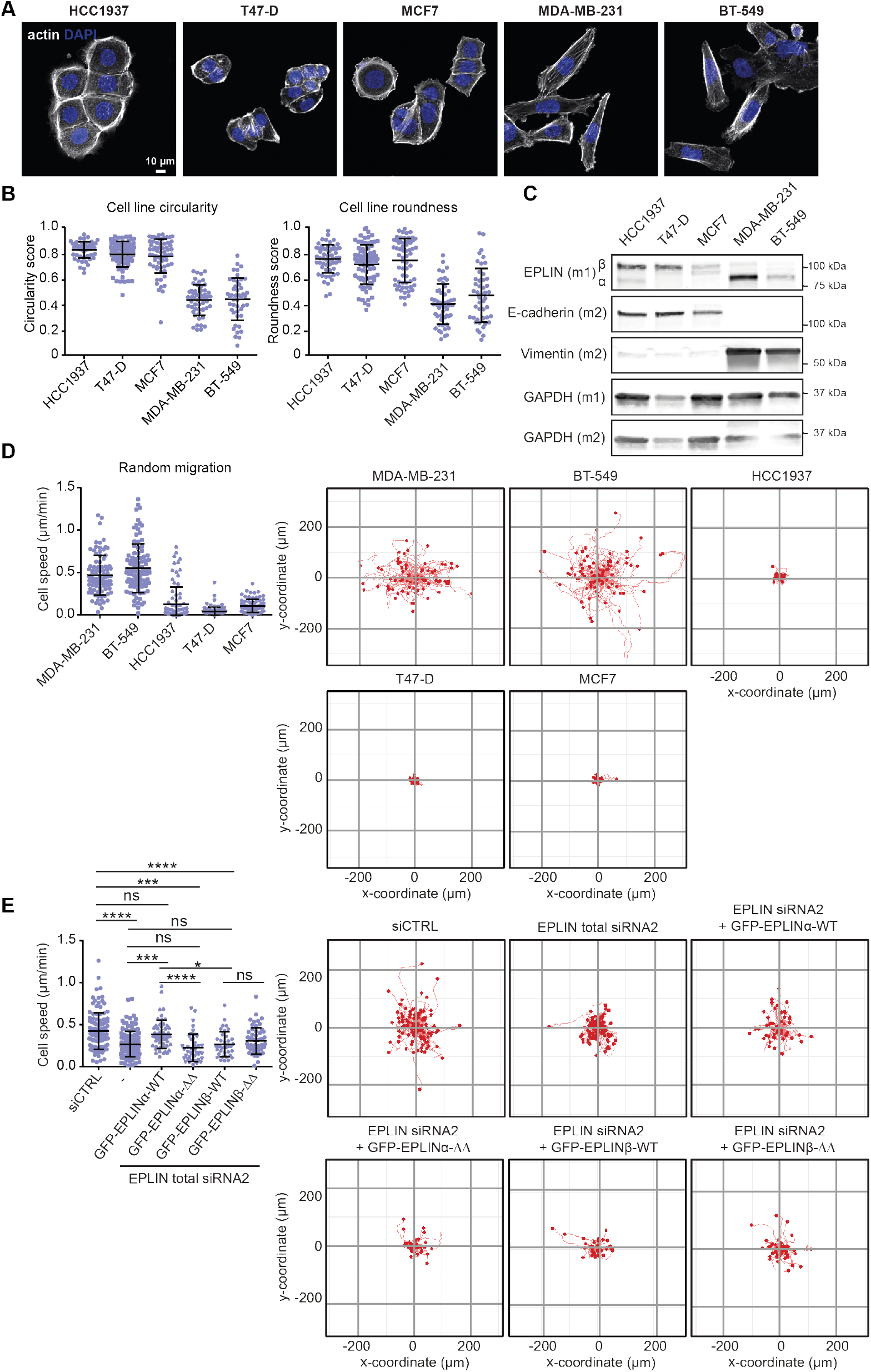
EPLINα mediates cell migration and correlates with a more mesenchymal phenotype in breast cancer cells. A) Representative confocal micrographs of the indicated cell lines labelled with Alexa Fluor 488 Phalloidin and DAPI. B) Cell circularity and cell roundness were analyzed for the indicated cell lines using Alexa Fluor 488 Phalloidin labelling to define the cell outline. HCC1937, n=57; T47-D, n=78; MCF7, n=66; MDA-MB-231, n=53; BT-549, n=46 analysed cells. C) Immunoblot showing expression of EPLIN isoforms, E-cadherin and vimentin in the indicated cell lines. m1=membrane 1, m2= membrane 2. Blots are representative of 3 independent experiments. D) Quantification of cell velocity of the indicated cell lines migrating randomly on plastic dishes. All migration tracks for each cell line pooled from three independent experiments are shown. MDA-MB-231, n=88 cells; BT-549, n=96 cells; HCC1937, n=87 cells; T47-D, n=89 cells; MCF7, n=74 cells. E) Quantification of cell velocity in siCTRL, EPLIN total siRNA2 or EPLIN total siRNA2 together with expression of GFP-EPLINα-WT / GFP-EPLINα- / GFP-EPLINβ-WT / GFP-EPLINβ- in MDA-MB-231 cells migrating randomly on plastic dishes. siCTRL, n=118 cells; EPLIN siRNA2, n=119 cells; siRNA2 + GFP-EPLINα-WT, n=55 cells; siRNA2 + GFP-EPLINα-, n=41 cells; siRNA2 + GFP-EPLINβ-WT, n=37 cells; siRNA2 + GFP-EPLINβ-, n=56 cells from three independent experiments. All migration tracks for each condition pooled from three independent experiments are shown. Plots from B, D, E) show mean±s.e.m. from three independent experiments. Statistical significance was assessed using Kruskal–Wallis one-way analysis of variance (ANOVA) and Dunn’s post hoc test. ****P<0.0001, ***P<0.001, *P<0.02, ns=not significant.

## Discussion

In this study, we report a novel, isoform-specific function of EPLINα in breast cancer. We show that the two EPLIN isoforms are localized to distinct subcellular actin-rich structures and are differentially expressed in epithelial versus mesenchymal cells. EPLINα is present on Rab21-and active β1-integrin-positive endosomes, localizes to tubules emanating from them and positively regulates integrin recycling to support cancer cell migration in an actin-binding dependent manner. Moreover, we determine the proximal interactomes of EPLIN isoforms in two cell lines, providing a rich resource for further exploration of the emerging isoform-specific roles of EPLINα and β. The current view on the role of EPLIN in cancer has remained unclear partly due to a gap in our knowledge regarding the expression pattern and function of EPLIN’s individual isoforms in different tissues. Our work emphasizes the importance of this balance and its implications in cancerous and healthy tissues, and our analysis of breast tumours highlights the relevance of EPLINα in triple negative breast cancer. Concordantly, Swip1, the cargo-adaptor of Rab21-mediated integrin endocytosis, is overexpressed in tumours from this molecular subtype (18); thus underlining the clinical impact of this specific active integrin transport pathway in breast cancer. Furthermore, recent work has demonstrated that EPLINα mediates migration of head and neck cancer cells (6, 7), which do not express EPLINβ. Taken together, our work identifies a unique role for EPLINα in the regulation of integrin trafficking and cell migration, and opens up new avenues to explore the roles of each EPLIN isoform in the context of breast cancer biology. We find that EPLINα binds to Rab21 directly. EPLINα also interacts directly with another Rab protein, Rab40, in a GTP-dependent manner (21). However, EPLINα is recruited to Rab40 via the SOCS domain, which is a unique domain of the Rab40 family and absent in Rab21 (44) implying that these Rab-proteins most likely interact and function with EPLIN via distinct molecular mechanisms. In our system, EPLINα is recruited predominantly to perinuclear Rab21-positive endosomes, perhaps due to the presence of F-actin hubs in these compartments(18). Thus, it is feasible to hypothesize that F-actin recruitment of EPLINα precedes Rab21 binding, and binding to Rab21 allows EPLINα to select the Rab21-associated integrin cargo for recycling. In liver hepatoma cells, EPLIN binds directly to NPC1L1 receptor at Rab11-positive endosomes, selecting it in this way for recycling (15, 45). It would be interesting to study whether EPLIN is recruited via F-actin in this context, and whether it can control recycling of other receptors. F-actin is required at all steps of endocytic transport (46–48). In the case of Rab21-mediated transport, F-actin is necessary for CLIC-mediated endocytosis, vesicle movement (18, 49), cargo sorting (50) and now we have identified a function for cargo recycling in an EPLINα-dependent fashion. The assembly of F-actin-related functional hubs at the endosomes seems to be cargo-specific. For example, WASH and retromer were shown to mediate the sorting of a subset of clathrin-independent cargos, such as MCT1, SLC3A2, Basigin and CD44 from Rab21-containing endosomes (50). Thus, the functional hubs assembled at Rab21-endosomes are able to recognize the cargo and the fate that they need to follow. Overall, our findings ratify the crucial role that F-actin plays at Rab21-containing endosomes. Here, we identified isoform-specific and shared interactors of EPLIN. Despite coronin 1C being a shared interactor of both isoforms, our results suggest that EPLINα is able to redistribute the cellular pool of coronin 1C by recruiting it to the endosomes. This would imply that the subcellular location of each EPLIN isoform can influence the function of the interactor. Hence, EPLINβ could have a more important role in cell-cell adhesion, or integrin adhesion stability, suggested by its preferential localization to adherens junctions and actin-stress fibers (Fig. 1C-D, Suppl. Fig. 3) (51). Furthermore, differences in actin-binding functions have been shown for each EPLIN isoform. In endothelial cells, EPLINβ stabilizes actin bundles during shear stress, while EPLINα prevents Arp2/3-mediated actin protrusions (20, 51). This is in accordance to our findings, where EPLINα is located at membrane protrusions. At the endosomes, this function could synergize with coronin 1C’s inhibition of actin branching by Arp2/3 to support tubule formation (37). Furthermore, coronin 1C has been shown to interact indirectly with EHD1, another member of FERARI that mediates endosomal fission (41). Thus, coronin 1C might have a dual role at recycling endosomes, restricting actin-branching and facilitating access of EHD1 to the tubulating endosomal membrane to catalyse fission. This conjecture is supported by the fact that EHD1 is required for integrin recycling (52). Our BioID approach also reflects the observations we made across the different cell lines and possible differential function of both EPLIN isoforms according to their localization. For example, some of the EPLINα-enriched interactors have a known link to endosomal transport, such as the phosphatase SYNJ1 and DPYSL2 (53, 54), whereas most of the identified EPLINβ-interactors have a role in stabilization of the actin cytoskeleton. For example, calponin isoforms CNN2 and CNN3 increase the formation and resistance of stress-fibers and filamin B and C crosslink actin, which also confers stability to stress fibers (32, 55). In addition, SHROOM3 localises to adherens juctions and rearrangements of the actin cytoskeleton to the apical side of epithelial cells (56). The functions of the EPLINβ-specific interactors are in accordance with the more epithelial-like phenotype observed in cells where this isoform is predominantly expressed, and might reflect an emerging role for EPLINβ in epithelial tissue homeostasis worth exploring in the future. Overall, our work reveals an isoform-specific function for EPLIN, highlights its impact on integrin recycling and cell migration in breast cancer cells and constitutes a valuable resource to explore its emerging shared and isoform-specific functions in normal and cancer cells.

## Materials and methods

### Lead contact

Further information and requests for resources and reagents should be directed to and will be fulfilled by the lead contact, Johanna Ivaska.

### Materials availability

All unique/stable reagents generated in this study are available from the lead contact with a completed materials transfer agreement.

### Data and code availability

All data that support the findings of this study are available within the paper and its Supplementary Information files. Any additional source data are available from the lead contact on request. This paper does not report original code.

### Cell lines

The human breast cancer cell lines MDA-MB-231, BT-549, HCC1937, T-47D and MCF7 were purchased from American Type Culture Collection (ATCC; cat. numbers in order: HTB-26, HTB-122, CRL-2336, HTB-133 and HTB-22). MDA-MB-231 cells were cultured in Dulbecco’s Modified Eagle’s Medium (DMEM) (VWR, 392-0413) supplemented with 10 % fetal bovine serum (Sigma, F7524), 2 mM L-glutamine (Sigma-Aldrich, G7513-100ML) and 1x non-essential amino acids (Sigma-Aldrich, M7145-100ML). BT-549 cells were cultured in RPMI-1640 medium (VWR, 392-0429) supplemented with 10 % fetal bovine serum (Sigma-Aldrich, F7524), 2 mM L-glutamine (Sigma-Aldrich, G7513-100ML) and 0.023 U/ml human insulin (Sigma-Aldrich, I9278-5ML). HCC1937 cells were cultured in RPMI-1640 medium (VWR, 392-0429) supplemented with 10 % fetal bovine serum (Sigma-Aldrich, F7524) and 2 mM L-glutamine (Sigma-Aldrich, G7513-100ML). T-47D cells were cultured in RPMI-1640 medium (VWR, 392-0429) supplemented with 10 % fetal bovine serum (Sigma-Aldrich, F7524), 2 mM L-glutamine (Sigma-Aldrich, G7513-100ML) and 0.2 U/ml human insulin (Sigma-Aldrich, I9278-5ML). MCF7 cells were cultured in DMEM (VWR, 392-0413) supplemented with 10 % fetal bovine serum (Sigma-Aldrich, F7524), 2 mM L-glutamine (Sigma-Aldrich, G7513-100ML) and 0.01 mg/ml human insulin (Sigma-Aldrich, I9278-5ML). U2OS cells were purchased from Leibniz Institute DSMZ-German Collection of Microorganisms and Cell Cultures and cultured in Dulbecco’s Modified Eagle’s Medium (DMEM) (VWR, 392-0413) supplemented with 10 % fetal bovine serum (Sigma, F7524) and 2 mM L-glutamine (Sigma-Aldrich, G7513-100ML). The cells were routinely tested for mycoplasma contamination and cultured at 37°C, 5 % CO2 in a humidified incubator.

### Transfection

The expression of proteins of interest was suppressed transiently using Lipofectamine RNAiMAX reagent (Thermo Fisher Scientific, 56532) according to the manufacturer’s instructions. The siRNAs were transfected at a concentration of 30 nM per oligo. The siRNA oligonucleotides targeting human LIMA1/Eplin were purchased from Horizon Discovery: Total Eplin siRNA1 (Accell A-010663-14, sequence GGCUUAAGAUGAUGUUUGA), Total Eplin siRNA2 (ON-TARGETplus J010633-09-0002, GCUUAAACAUUACGACUGA) and Eplin β siRNA1 (Dharmacon_A010663-13, UCACUAUCAUUGAGGGUAA). Eplin β siRNA2 was designed using the online tool BLOCK-iT™ RNAi Designer and custom-made from Invitrogen, (CAGGUAUUUCAGUGUCUGUAGA-CAA). The siRNA used as control (siCTRL) was Allstars negative control siRNA (Qiagen, cat. no. 1027281). Plasmids of interest were transfected using Lipofectamine 3000 (Invitrogen, 100022052) according to the manufacturer’s instructions.

### Plasmids

Eplin α-GFP and Eplin β-GFP were obtained from Addgene (plasmids 40947 and 40948, respectively). To generate the mScarlet-I-tagged Eplin α, PCR was first performed using pEFHD2-mScarlet-I as a template for mScarlet-I with the mScarlet_F GATCCGCTAGCGC-TACCGGTCGCCACCATGGTGAGCA and mScarlet_R GGATCTGAGTCCGGACTTGTACAGCTCGTCCATGCC primers. Overlap extension PCR was then performed for Eplin α using the pEGFP-Eplinα (Addgene plasmid 40947) as template and the Eplin_F ACGAGCTGTACAAGTC-CGGACTCAGA and Eplin_R CCGGTGGATCCCGGGC primers. The overlapped fragments were then digested with NheI/KpnI (New England Biosciences) restriction enzymes and ligated into the similarly digested pEGFP-Eplinα backbone using T4 DNA Ligase (ThermoFisher, EL0011). All plasmids were validated by analytical digests and sequencing. In order to generate the BioID constructs, the entry vectors pENTR2B-EPLINα and pENTR2B-EPLINβ were first generated using HiFi DNA assembly according to the manufacturer’s instructions (New England Biolabs).The primer sequences used to make the plasmids are shown below. Vectors were linearised using restriction enzymes indicated for 1 hour at 37°C. PCR-amplified inserts and linearised vectors were purified (and sizes confirmed) using gel extraction. To generate pPB-TagBFP-T2A-myc-BirA* (pPB-BirA*), TagBFP-T2A-myc-BirA* was amplified from pCDH-TagBFP-myc-BirA* (57, 58) and inserted into a pPB destination vector, pPB-DEST (59), linearized by BspDI and XhoI. To generate EPLINα and EPLINβ BioID plasmids (pPB-TagBFP-T2A-myc-BirA*-EPLINα and pPB-TagBFP-T2A-myc-BirA*-EPLINβ; herein referred to as BirA*-EPLINα and BirA*-EPLINβ, respectively), full length EPLIN ORFs (Addgene plasmids 40947 and 40948) were amplified by PCR and inserted into the pPB-TagBFP-T2A-myc-BirA* vector that was linearised using BspDI and XhoI. For pENTR2B-EPLINα, full length EPLIN ORFs were amplified by PCR and inserted into the pENTR2B vector (Gateway™ pENTR™ 2B Dual Selection Vector, Invitrogen) linearised by SalI and EcoRV restriction enzymes. pDEST_V2-EPLINα was generated by gateway cloning. Using LR Clonase II (Invitrogen), pENTR2B-EPLINα was LR subcloned with a pDEST-V2-ORF (Addgene 73636) according the manufacturers’ protocol. Sequences were confirmed using DNA sequencing. pLL5.0-hCORO1C-mEmerald was a gift from James Bear (University of North Carolina-Chapel Hill School of Medicine, Chapel Hill, North Carolina, USA). StrepII-EPLINα WT plasmid was created by inserting the EPLINα PCR product (created with EPLINα Strep F/R primers) into pcDNA3 StrepII MCS vector cleaved with EcoRI/AgeI using NEBuilder HiFi DNA Assembly (NEB). The deletion mutants (ΔNHX, ΔWH2) were created using whole plasmid synthesis approach using the primers specified below. Briefly, after PCR with the indicated primers and StrepII-EPLINα WT pcDNA3 as a template, the reaction has been cleaned using KAPA Pure Beads (Roche), digested with DpnI and transformed into

### Primers used to contruct the EPLIN plasmids

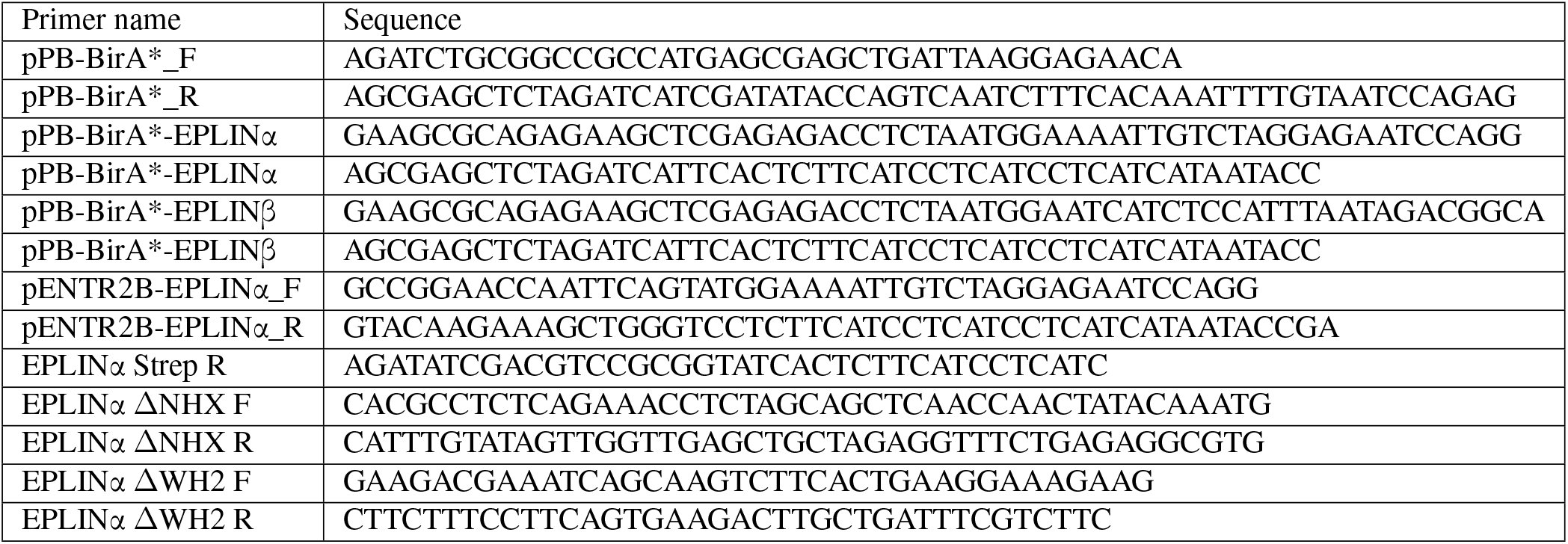

DH5α E. coli strain. The actin binding-deficient mutant StrepII-EPLINα ΔΔ pcDNA3 was generated introducing the ΔNHX mutation into StrepII-EPLINα ΔWH2 pcDNA3 plasmid. The mScarlet-I/GFP EPLINα mutant constructs were generated as above using mScarlet-I/GFP EPLINα WT as template. All the constructs were verified by sequencing.

### Western blotting

Cells were washed with cold PBS and scraped into a TX lysis buffer (TXLB; 50 mM Tris-HCl, pH 7.5, 150 mM NaCl, 0.5% Triton-X, 0.5% glycerol, 1% SDS, Complete protease inhibitor (Sigma-Aldrich, 5056489001), and phos-stop tablet (Sigma-Aldrich, 4906837001). SDS sample buffer was added to the lysates, samples were placed on a heat block (+90°C) for 10 minutes and separated by SDS–polyacrylamide gel electrophoresis (4–20% Mini-PROTEAN TGX gels, Bio-Rad, 456-1096). Proteins were transferred to a nitrocellulose membrane (Bio-Rad, 1704159) using the Trans-Blot Turbo transfer system (Bio-Rad) and the membrane was blocked with StartingBlock blocking buffer (Thermo Fisher Scientific, 37538) for 1 hour at room temperature. Primary antibodies against proteins of interest were added to the membrane and the membrane was incubated with the antibody solution overnight at +4°C, followed by incubation with fluorophore-conjugated secondary antibodies for 1 hour at room temperature. All antibodies were diluted in StartingBlock blocking buffer (Thermo Fisher Scientific, 37538). Membranes were scanned with the Odyssey infrared imaging system (LI-COR Biosciences).

### BiFC microscopy

Cells plated on 6-well plates were co-transfected with split Venus constructs pDEST-V2-EPLINα and pDEST-Rab21-V1 (18). The next morning, the transfected cells were detached using trypsin, and plated on glass-bottom dishes (Cellvis, D35-14-1.5-N). Samples were fixed 5 hours after plating with warm 4% paraformaldehyde for 15 minutes. The above-mentioned steps for immunofluorescence staining were then followed. The samples were imaged with a 3i (Intelligent Imaging Innovations, 3i Inc) Marianas spinning disk confocal microscope, Hamamatsu ORCA-Flash4.0 v2 scientific complementary metal-oxide semiconductor (sCMOS) camera (Hamamatsu Photonics), and Plan-Apochromat 63x, 1.4 NA oil objective. Analysis of colocalization was performed with the Coloc2 plug-in for Fiji (ImageJ, National Institutes of Health).

### Analysis of cell circularity and roundness

Cell circularity and roundness of cells were analyzed using Fiji (ImageJ, National Institutes of Health). Cell outlines were drawn based on F-actin staining, and circularity and roundness were selected as measurement parameters from Fiji’s built-in shape descriptors function before performing the measurement. Fiji uses the formula 4π × [Area]/[Perimeter]2 to calculate circularity and the formula 4 × [Area]/π × [Major axis]2 to calculate roundness.

### Migration assay

Cells were plated on a 6-well plate in full growth medium. Before imaging, medium was changed to full growth medium containing 25 mM HEPES. Cells were imaged every 10 or 20 minutes for 6 hours at +37°C and 5% CO2 using a Nikon Eclipse Ti-E epifluorescence microscope and Plan Fluor 10x, 0.30 NA objective (Nikon), controlled by NIS-Elements AR 4.60 software (Nikon). To quantify cell migration, single migrating cells were tracked with the MTrackJ plug-in for Fiji (ImageJ, National Institutes of Health). Cells were tracked based on phase contrast signal or, in the case of cells expressing a plasmid construct, based on fluorescent protein signal. Dividing or dying cells were excluded from the analysis.

### Immunofluorescence staining, SIM, Airyscan microscopy and live imaging

Cells plated on glass-bottom dishes (Cellvis, D35-14-1.5-N or MatTek, P35G-0.170-14-C) were fixed with 2% paraformaldehyde for 15 minutes, permeabilized for 15 minutes with 0.3% Triton X-100 in 10% horse serum (Gibco, 16050-122) in PBS and blocked for 20 minutes with 10% horse serum in PBS. Samples were incubated with primary antibodies diluted in 10% horse serum overnight at +4°C. Secondary antibodies were diluted in 10% horse serum and the samples were incubated with the secondary antibodies for 1-2 hours at room temperature. Where indicated, cell nuclei were stained using DAPI and filamentous actin with Phalloidin-Atto 647N (Sigma-Aldrich, 65906), Alexa Fluor 488 Phalloidin (Invitrogen, A12379) or SiR-actin (Spirochrome, SC001). For SIM and Airyscan imaging, cells plated on glass-bottom dishes (MatTek, P35G-0.170-14-C) were fixed with 4% paraformaldehyde for 15 minutes and treated as described above, with an additional post-fixation step (4% paraformaldehyde, 10 min) after incubation with secondary antibodies and Phalloidin-Atto 647N. Structured illumination microscopy (SIM) was performed using a DeltaVision OMX v4 (GE Healthcare Life Sciences) with a front-illuminated pco.edge sCMOS camera (pixel size 6.5 mm, readout speed 95 MHz; PCO AG) and a Plan-Apochromat 60x, 1.42 NA objective (immersion oil RI of 1.516). SIM illumination mode (five phases x three rotations) was used, and the microscope was controlled by SoftWorx. Airyscan imaging was performed using a LSM880 laser scanning confocal microscope (Zeiss) equipped with an Airyscan detector and 63x oil, NA 1.4 objective. Standard super-resolution mode was used for Airyscan imaging, and the microscope was controlled with Zen Black (2.3). Live imaging of tubulating endosomes was performed using a Leica Stellaris 8 Falcon FLIM microscope equipped with a Plan APO 63x 1.40 NA oil immersion objective. Acquisition was done for 252 frames every 0.66 seconds using a pinhole setting of 1AU at 580 nm with unidirectional scanning with the resonance scanner at a line speed of 8kHz. Detection was done in photon counting mode with a line accumulation of 10 lines. Image channels were in Track 1, simultaneous excitation at 487 nm and 650 nm with detection windows at 488-530 nm (HyD X detector) and 658-717 nm (HyD X detector), and in Track 2 excitation at 569 nm and emission window of 582-640 nm (HyD S detector).

### Endogenous tagging of Rab21

MDA-MB-231 cells expressing endogenously tagged Rab21 were generated using homology-directed repair-mediated (HDR) CRISPR/Cas9 targeting Exon 1 of Rab21-201 transcript (Transcript ID ENST00000261263.5). Corresponding guide sequence (AGCGACGGGATGGCTGCGGC) was cloned into pXPR_001 lentiCRISPR v1 plasmid (Addgene 49535) cleaved with BsmBI (NEB). Final construct (pXPR_001-sgRAB21) was verified by sequencing. The template for HDR was designed based on Zhang et. al., 2017 (60) and synthesized into pUC57 mini plasmid (final plasmid pUC57-RAB21 HDR; Genscript). The sequence of mScarlet-I, followed by a short linker, was surrounded by approximately 850 bp-long homology arms and flanked with Rab21 Cas9 guide sequence to generate double-cut dsDNA donor sequence for recombination. MDA-MB-231 cells were co-transfected with pXPR_001-sgRab21 and pUC57-Rab21 HDR plasmids in 1:4 ratio using Lipofectamine 3000 (Invitrogen). Five days post-transfection, the cells were sorted using SONY SH800S Cell Sorter for red fluorescence, expanded, and resorted again to obtain the cell population with the highest fluorescence intensity.

### Integrin internalization assay

Cells on Cellvis glass-bottom dishes were placed in +4°C to cool down for 15 minutes, after which they were incubated on ice with primary antibody against active integrin β1 (12G10) diluted in cold full growth medium for 1 hour to label cell surface integrins. Samples were washed twice with cold full growth medium, and warm full growth medium was added before placing the samples in an incubator at +37°C to allow antibody-labeled integrin internalization for 15 minutes. Internalization was then stopped by changing to cold medium, and samples were placed on ice. Samples were washed twice with an acid wash solution (0.2M acetic acid and 0.5M NaCl, pH2.5) to remove the remaining antibodies from the cell surface. Finally, samples were fixed with 2% paraformaldehyde for 20 minutes at room temperature. Quantification of internalized integrins was performed on 3D projections of the cells using the ‘spots detection’ function of IMARIS software. The intensity sum of all of the vesicles in a cell was divided by the cell’s volume. All intensity values were then normalized to the average of the control condition (siCTRL).

### Integrin recycling assay

Cells were starved for 45 minutes in warm serum-free medium at +37°C, after which samples were cooled down at +4°C for 10 minutes before fluorophore-conjugated 12G10 (12G10 conjugated to Alexa 488, 12G10-488), diluted in cold serum-free medium, was added to the samples and incubated for 1 hour on ice to label cell surface integrins. Samples were washed twice with cold serum-free medium, after which warm serum-free medium was added before placing the samples in an incubator at +37°C to allow antibody-labeled integrin endocytosis for 30 minutes. Endocytosis was then stopped by adding cold serum-free medium and placing samples on ice. Next, an anti-Alexa 488 antibody, diluted in cold serum-free medium (50 µg/ml), was added to the samples and incubated for 1 hour on ice. The anti-Alexa 488 antibody binds to 12G10-488-bound integrins, effectively quenching the fluorescent signal from the integrins left on the cell surface. To start integrin recycling, warm full growth medium containing the anti-Alexa 488 quenching antibody was added to the samples and incubated at +37°C for 30 minutes. As 12G10-488-bound integrins are recycled from inside the cell and reach the cell surface, their fluorescence is effectively extinguished by the quenching antibody in the medium. Finally, samples were fixed with 2% paraformaldehyde for 20 minutes at room temperature. Samples were stained with Phalloidin-Atto 647N (see above) and imaged with a 3i (Intelligent Imaging Innovations, 3i Inc) Marianas spinning disk confocal microscope with a Yokogawa CSU-W1 scanner and a back-illuminated 10MHz EMCDD camera (Photometrics Evolve) with a 63x/1.4 oil objective. Intracellular integrin signal intensity per cell, normalized to cell area, was quantified using Fiji (ImageJ, National Institutes of Health) using mean background-subtracted sum projections of 21 slices from the middle of the cell.

### Microscale thermophoresis

For microscale thermophoresis (MST), GppNHp-loaded Rab21 (18) was labeled using amine-reactive NT-650-NHS fluorescent dye (Nanotemper) using 1:2 protein:dye concentration ratio. Excess dye was removed by centrifugation using centrifugal filters (Millipore). GST-tagged EPLINα was expressed in E. coli and purified as described elsewhere. Protein concentrations were determined using Bradford assay (Bio-Rad). Binding experiments were carried out in 10 mM HEPES, pH 7.4, 150 mM NaCl, 5 mM MgCl2, 0.05% Tween-20. The binding assays were performed using a fixed concentration (50 nM) of the fluorescently labeled Rab21(target) and two-fold serially diluted decreasing concentrations of the unlabeled EPLINα (ligand). 12 serial dilutions of the ligand and labelled protein were loaded into the standard capillaries (Nanotemper, MO-K022) and analyzed using the Monolith NT.Automated instrument. All experiments were carried out at RT using a medium MST power. Normalization of the fluorescence signal and fitting to the KD equation were performed using the software MO Affinity Analysis v1.6 (Nanotemper).

### Endosome line scan analysis

For line scan analyses of endosomes, a single-plane image from the middle of the endosome was used. A circular, clockwise freehand line was drawn around the endosome, after which intensity values of the different channels were measured, normalized to values between 0-100, and plotted as a function of length.

### Analysis of coronin 1C signal after EPLIN knockdown

MDA-MB-231 cells expressing endogenously tagged Rab21-mScarlet and plated on glass-bottom dishes (MatTek, P35G-0.170-14-C) were fixed and immunostained as described earlier. Samples were imaged using a LSM880 laser scanning confocal microscope (Zeiss) equipped with a 63x oil, NA 1.4 objective and controlled with Zen Black (2.3) software. The Rab21-mScarlet signal was used to find a suitable single z-plane dominated by endosomal Rab21 signal, and a cytoplasmic region of interest was drawn based on the Rab21-mScarlet signal. Then, signal intensity of the coronin 1c immunofluorescence staining was measured after background subtraction.

### Deconvolution

Where indicated, AiryScan images were deconvolved using Huygens Professional version 24.04 (Scientific Volume Imaging). Raw data were analysed using Array Detector Quality Control tool and deconvolved using CMLE algorithm with 15 iterations and the recommended settings.

### BioID and affinity purification of proximity-biotinylated proteins

For stable expression of pPB-BioID constructs (BirA*, BirA*-EPLINα and BirA*-EPLINβ) in 231 and HCC1957 cells, pPB-BioID constructs were co-transfected with a plasmid containing the piggyBac transposase, pCMV hypBase (61), with a 3 to 1 ratio (pPB-BioID vector : pCMV-HypBase) using Lipofectamine 3000 according to manufacturers’ instructions. Fresh media was added to cells the following day. Cells were passaged for two weeks before being FACs sorted for BFP expression to select high- and low-expressing cells. Cell populations were selected based on expression levels and subcellular targeting of BirA*-tagged proteins, aiming for equal levels of expression and efficient subcellular targeting. Although the low-expressing BirA*-EPLINα and BirA*-EPLINβ and high-expressing BirA* control cells were selected, BirA* expression levels were significantly lower. For biotinylation of proximal proteins, cells were seeded onto 10 cm dishes (two per condition) one day before the addition of biotin (from 20X solution in media, final concentration of 50 µM) to induce biotinylation. After 24h incubation with biotin, cells were lysed and biotinylated proteins affinity-purified following a previously described BioID protocol (57, 62) adapted from Roux et al., 2012-2013 (63, 64). At RT, cells were washed 3 x with 10 ml PBS before 400 µl cell lysis buffer added (250 mM NaCl, 50 mM Tris HCl (pH 7.4), 0.1 % SDS (w/v), 0.5 mM DTT and 1X protease inhibitors (cOmplete Mini, EDTA-free, Roche). Cells were scraped, and lysates (800 µl per condition) added to 80 µl 20% (v/v) Triton X-100. From herein, lysates were kept at 4°C. Lysates were passed four times through a 19G needle before 720 µl 50 mM Tris HCl (pH 7.4) added. Lysates were then passed through a 27G needed a further 4 times, before centrifugation for 10 m at full speed at 4°C. The supernatant was incubated with 30 µl MagReSyn magnetic streptavidin beads (ReSyn Biosciences; washed twice with lysis buffer) overnight at 4°C, with rotation. The following day, the beads were washed with multiple stringent buffers (500 µl each, 5 min) to remove non-specific proteins: twice with wash buffer 1 (10% SDS (w/v)), followed by one wash each of wash buffer 2 (500 mM NaCl, 50 mM HEPES, 1 mM EDTA, 1% Triton X-100 (w/v), 0.1% deoxycholic acid (w/v)) and wash buffer 3 (10 mM Tris HCl (pH 7.4), 1 mM EDTA, 0.5 % NP-40 (w/v), 0.5% deoxycholic acid (w/v)). To elute proteins, beads were incubated with 90 µl 2X reducing sample buffer with 10 µM biotin at 70°C for ten minutes. Western blotting confirmed the presence of biotinylated proteins before the samples prepared for label free quantitative mass spectrometry (MS).

### MS sample preparation

Samples were prepared for MS using in-gel trypsin digestion. Samples were subjected to SDS-PAGE for 4 minutes (or until all the sample had entered the gel) at 200V (two wells per sample; 4-20% Mini-PROTEAN TGX precast protein gel, BIO-RAD), then stained with Coomassie blue for ten minutes before being washed 4 X with ddH2O (10 minutes each, with a final wash overnight at 4°C). Protein bands were excised and gel pieces were washed twice with 0.04 M NH4HCO3/50% acetonitrile (ACN) for 15 mins, before incubation with 100% ACN for 10 mins. Proteins were reduced with 20 mM DL-Dithiothreitol (DTT; BioUltra, for molecular biology, Sigma) at 56°C for 30 minutes, followed by addition of 100% ACN for 10 mins, and proteins alkylated with 55 mM iodoacetamide (Sigma) in 100 mM NH4HCO3 for 20 mins at RT (protected from light). Gel pieces were washed twice with 100 mM NH4HCO3 and dehydrated with 100% ACN followed by centrifugation in a vacuum centrifuge. Proteins were digested using 0.005 µg/µl trypsin (sequencing grade modified trypsin; Promega) in 40 mM NH4HCO3/10% ACN at 4°C for 20 minutes followed by 16 h at 37°C. Peptides were extracted by incubating with 100% ACN followed by 50% ACN/5% formic acid (Thermo Fisher) for 15 mins each at 37°C. The supernatant was collected after each step of peptide extraction, and dried in a vacuum centrifuge. Peptides were dissolved in 10 µl 2% formic acid immediately prior to MS analysis.

### MS data acquisition

Samples were analysed by liquid chromatography-electrospray ionisation-tandem mass spectrometry (LC-ESI-MS/MS) using a nanoflow HPLC system (Easy-nLC1200, Thermo Fisher Scientific) coupled to a Q Exactive HF mass spectrometer (Thermo Fisher Scientific, Bremen, Germany) equipped with a nano-electrospray ionization source. Peptides were first loaded on a trapping column and subsequently separated inline on a 15 cm C18 column (75 µm x 15 cm, ReproSilPur 3 µm 120 Å C18-AQ, Dr. Maisch HPLC GmbH, Ammerbuch-Entringen, Germany). The mobile phase consisted of water with 0.1% formic acid (solvent A) and ACN/water (80:20; v/v) with 0.1% formic acid (solvent B). To elute peptides, a 30 min linear gradient from 6% to 39% of solvent B was used, followed by a wash stage with 100% eluent B. MS data were automatically acquired using Thermo Xcalibur 4.1 software (Thermo Fisher Scientific). Data dependent acquisition was used, consisting of repeated cycles of a single MS1 scan covering a range of m/z 350 – 1750 plus a series of HCD fragment ion scans (MS2 scans) for up to 10 of the highest intensity peptide ions from the MS1 scan.

### MaxQuant processing and bioinformatic analyses

MS data were processed and bioinformatic analyses performed as described previously (57, 62). Raw MS data were analysed using MaxQuant (v2.0.3.0, available from Max Planck Institute of Biochemistry) (65, 66) using default parameters, with biotinylation of lysines as a variable modification and selecting match between runs, LFQ quantification, and unique peptides only for quantification. Experiments were searched against the human proteome (UniProtKB/Swiss-Prot (67) accessed April 2023). SAINT-express (28) (via REPRINT; https://reprint-apms.org/) was used to identify high-confidence bait-prey interactions using LFQ intensities from MaxQuant, using default parameters and MS1-based processing. A BFDR of 0.05 was used as a threshold to identify bait-prey interactions considered as proximal interactors. Dotplots were generated using R and prey organised by hierarchical clustering (Jaccard distance). Prey groups were assigned numbers (1-12) to aid identification of prey. Gene ontology analysis was performed using ClusterProfiler (version 4.8.3) (39). Interactome networks were generated using Cytoscape, version 3.10.1 (69). Actin-binding proteins were identified using UniProt Annotated Keywords via STRING (70). ‘Known EPLIN interactors’ were those identified from BioGRID, accessed February 2024 (71).

### Breast tumour sample analysis

The breast cancer patient cohort has been previously described in Milde-Langosch et. al., 2014 (72). Briefly, tumour samples were obtained from 105 primary breast cancer patients treated at the University Medical Center Hamburg Eppendorf, Germany, Department of Gynecology between 1991-2002. All tissue samples were snap-frozen after surgery and stored in liquid nitrogen until use. From those samples, vimentin mRNA levels were analysed using microarray data (Affymetrix, Santa Clara, CA, USA) after RNA extraction and statistical analyses were conducted using SPSS software Version 23 (SPSS Inc., Chicago, IL, USA). Protein extracts from the same samples were obtained by lysing tumour fragments with RIPA buffer to analyse EPLIN isoforms expression. Tumour lysates were subjected to SDS-PAGE followed by protein transfer and immunoblotting with primary antibodies (anti EPLIN, Novus Cat NB100-2305 and anti β-actin, Santa Cruz Cat sc-47778). Signal amplification was performed with HRP-conjugated secondary antibodies (Goat Anti Mouse IgG and Anti Rabbit IgG von Southern Biotek, 130-05 1:8000, Lot: E2518-Z909), diluted 1:8,000 in blocking buffer, at room temperature for 1h. The membranes were developed and expression of each EPLIN isoform was quantified using ImageJ. The ratio between both isoforms was calculated (EPLINβ/EPLINα). The cohort was then divided into four groups according to the ratio of EPLIN isoforms (cutoff values: Group β>α, ratio >1.3, Group α>β, ratio <0.7, Group α=β, ratio between 0.7 and 1.3) and the clinicopathological factors (ER status and molecular subtype) of each sample correlated and plotted as the number of patients.

### Statistical methods

Statistical analyses and plotting of data were performed using Prism 7 (GraphPad). P values and statistical tests used are indicated in the figure legends. RStudio was used to plot migration tracks.

## Acknowledgements

We thank P. Laasola and J. Siivonen for technical assistance, and the Ivaska laboratory for critical reading of the manuscript and constructive feedback over the course of this study. We thank James Bear (University of North Carolina-Chapel Hill School of Medicine, Chapel Hill, North Carolina, USA) for providing reagents. The Cell Imaging and Cytometry core (Turku Bioscience Centre, University of Turku and Åbo Akademi University and Biocenter Finland) are acknowledged for services, instrumentation and expertise. We also thank Christoffer Lagerholm for assistance with image acquisition. This work was supported by the Finnish Cancer Institute (K. Albin Johansson Professorship, J.I.); a Research Council of Finland project grant (# 325464 to J.I.) and Centre of Excellence program (# 346131, J.I.); the Cancer Foundation Finland (J.I.); the Sigrid Juselius Foundation (J.I.); the Research Council of Finland’s Flagship InFLAMES (# 337530 & 357910) and the Jane and Aatos Erkko Foundation (J.I.). N.Z.J. was supported by the University of Turku Doctoral Programme in Technology, the Swedish Cultural Foundation in Finland, the Varsinais-Suomi Regional Fund, the K. Albin Johansson Foundation and the Ida Montin Foundation. This study has been supported by the Research Council of Finland postdoctoral fellowships (grant nos. 321493 (to P.M.-L.) and grant 338585 (to J.R.W.C)). J.R.W.C. was also supported by the European Union’s Horizon 2020 research and innovation programme under the Marie Sklodowska-Curie grant agreement [grant 841973].

## Author contributions

Conceptualization: N.Z.J., P.M.-L. and J.I. Methodology: N.Z.J., P.M.-L. and J.I. Formal analysis and investigation: N.Z.J., P.M.-L., M.R.C., M.D., J.R.W.C., V.-M.L., K.E., L.O.-F., S.V. and J.I. Resources: S.V., L.O.-F. and K.E. Writing–original draft: N.Z.J., P.M.-L., M.R.C., M.D., J.R.W.C. and J.I. Writing–review and editing: N.Z.J., P.M.-L., M.R.C., M.D., H.H., J.R.W.C. and J.I. Visualization: N.Z.J., P.M.-L., M.R.C., M.D., H.H. and J.I. Supervision: P.M.-L. and J.I. Funding acquisition: N.Z.J., P.M.-L., and J.I.

