## Supplementary information for "EPLINα controls integrin recycling from Rab21 endosomes to drive breast cancer cell migration"

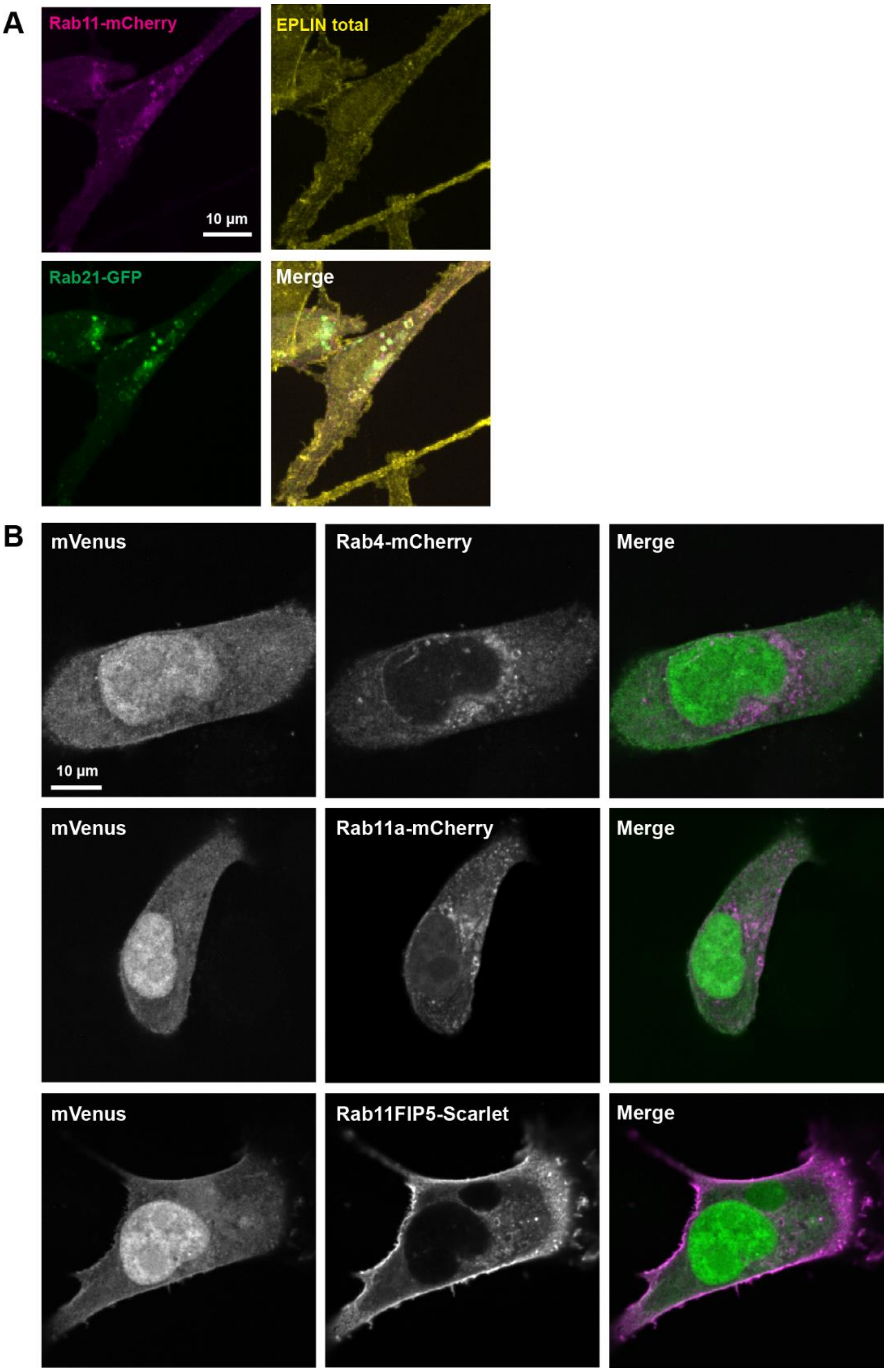


**Suppl. Fig1.** A) Representative confocal images of MDA-MB-231 cells transfected with Rab21-GFP and Rab11-mcherry, and stained for total EPLIN. B) Representative images of MDA-MB-231 cells transfected with mVenus control together with proteins fused to mcherry/Scarlet. These cells were the negative control for experiments in Figure 2D.

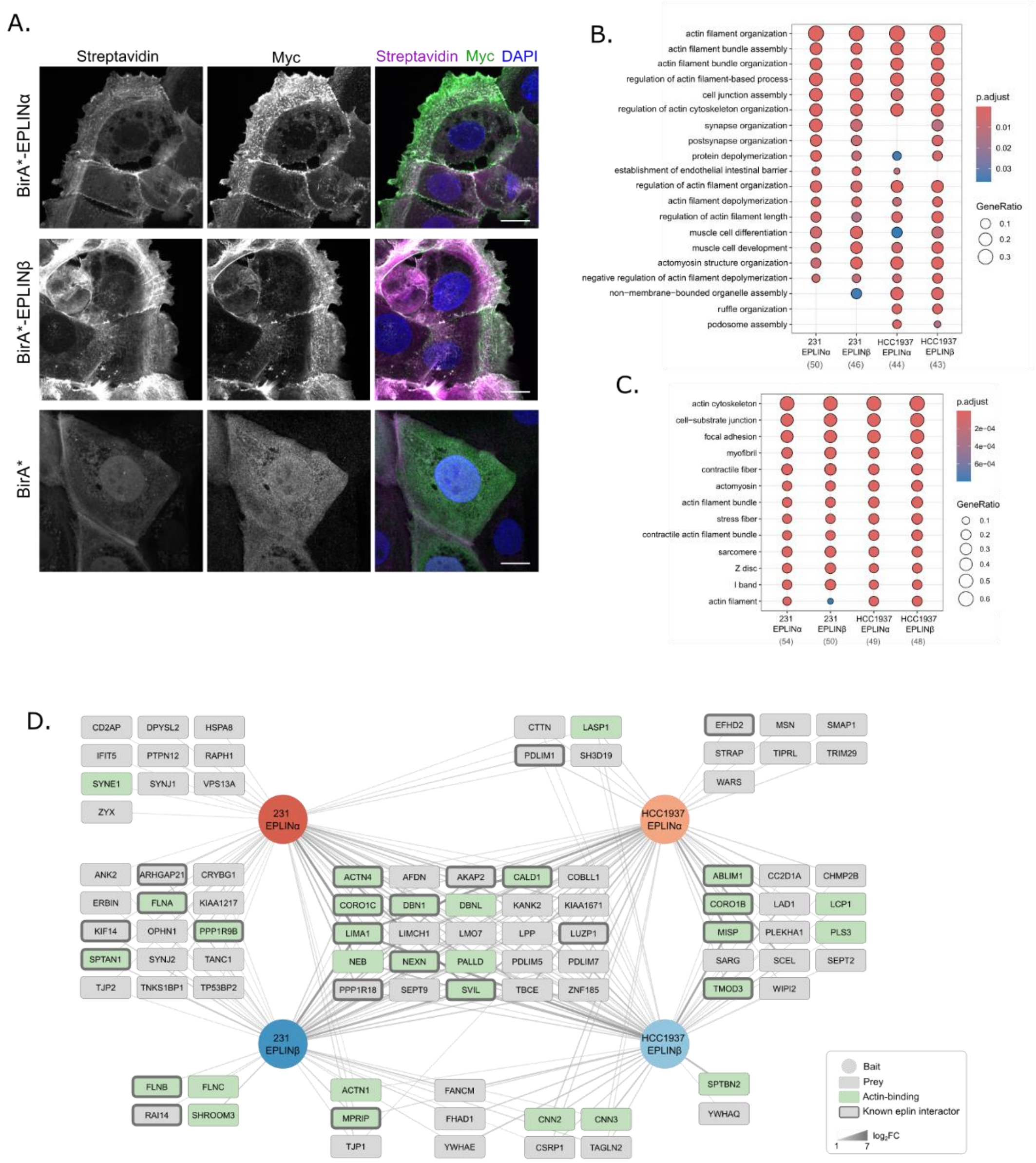


**Suppl. Fig2.** A) Immunofluorescence images of HCC1937 cells stably expressing BirA*-tagged EPLINα, EPLINβ or BirA* control. Cells were incubated with biotin for 24 hours to induce labelling, and biotinylated proteins detected by fluorescently-conjugated streptavidin, and BirA*-tagged proteins detected by anti-myc antibody and nuclei were stained with DAPI. B, C) Enriched gene ontology terms of high-confidence proximity interactors (BFDR ≤ 0.05) of BirA*-EPLINα and BirA*-EPLINβ identified by BioID in 231 and HCC1937 cells in the Cellular Component (C) and Biological Process categories. Number of gene names used for analysis indicated in brackets. p.adjust; adjusted p value, GeneRatio; ratio of genes identified in terms. GO analysis performed using ClusterProfiler (Wu et al., 2021). D) Proximity interaction network of BirA*-EPLINα and BirA*-EPLINβ in 231 and HCC1937 cells. Circular nodes indicate baits (BirA*-EPLINα/β), and rectangular nodes indicate prey (proximity interactors, BFDR ≤ 0.05). Previously identified EPLIN interactors (BioGRID) are indicated by thick grey borders, and actin-binding (UniProt Annotated Keywords) are shown in green. Edge width and colour indicate fold change over BirA* control (log2FC).

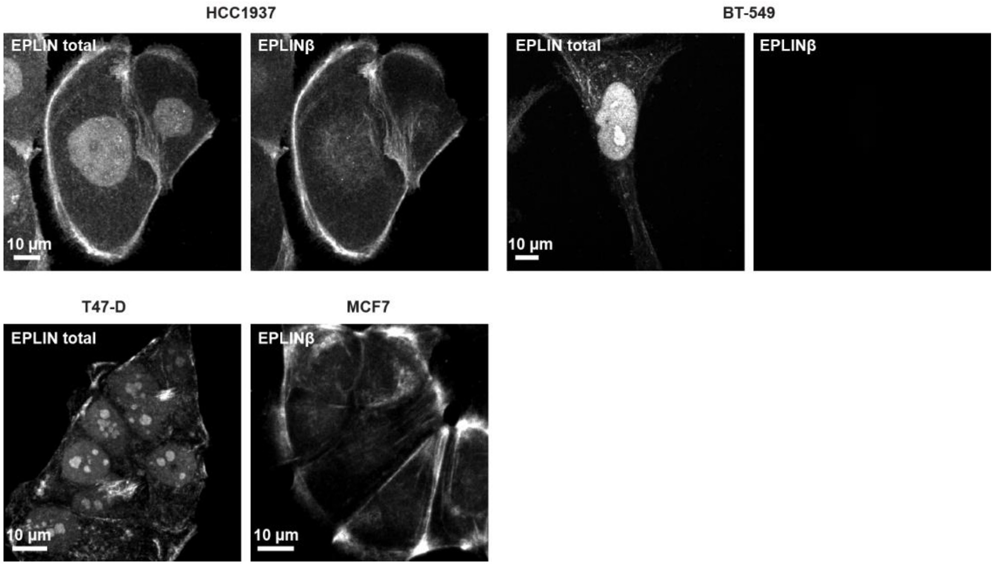


**Suppl. Fig3.** Representative confocal images of EPLIN total and/or EPLINβ immunostaining (using Eplin monoclonal antibody OTI1A3, Thermo Fisher MA5-27016) in HCC1937, BT-549, T47-D and MCF7 cells.

### Supplementary Videos

Suppl. video 1. mScarlet-EPLINα localizes at tubules emerging from Rab21-containing endosomes. MDA-MB-231 live cells stably expressing Rab21-GFP and mScarlet-EPLINα. Related to Figure 2F.

Suppl. video 2. Localization of EPLINα to endosomes requires an intact actin-binding function. MDA-MB-231 live cells overexpressing mScarlet-EPLINα-WT and GFP-EPLINα-ΔΔ (actin-defective binding). Related to Figure 3F.

Suppl. video 3. Random migration of MDA-MB-231 cells, related to Figure 7D.

Suppl. video 4. Random migration of BT-549 cells, related to Figure 7D.

Suppl. video 5. Random migration of HCC1937 cells, related to Figure 7D.

Suppl. video 6. Random migration of T47-D cells, related to Figure 7D.

Suppl. video 7. Random migration of MCF7 cells, related to Figure 7D.

Suppl. video 8. Random migration of siCTRL-transfected MDA-MB-231 cells, related to Figure 7E.

Suppl. video 9. Random migration of EPLIN total siRNA-transfected MDA-MB-231 cells, related to Figure 7E.

Suppl. video 10. Random migration of EPLIN total siRNA-transfected MDA-MB-231 cells expressing GFP-EPLINα-WT, related to Figure 7E.

Suppl. video 11. Random migration of EPLIN total siRNA-transfected MDA-MB-231 cells expressing GFP-EPLINα-ΔΔ, related to Figure 7E.

Suppl. video 12. Random migration of EPLIN total siRNA-transfected MDA-MB-231 cells expressing GFP-EPLINβ-WT, related to Figure 7E.

Suppl. video 13. Random migration of EPLIN total siRNA-transfected MDA-MB-231 cells expressing GFP-EPLINβ-ΔΔ, related to Figure 7E.

**Supplementary Table**

Table 1. Proximity interactomes of EPLINα and EPLINβ.
